# The statistics of *k*-mers from a sequence undergoing a simple mutation process without spurious matches

**DOI:** 10.1101/2021.01.15.426881

**Authors:** Antonio Blanca, Robert S. Harris, David Koslicki, Paul Medvedev

## Abstract

K-mer-based methods are widely used in bioinformatics, but there are many gaps in our understanding of their statistical properties. Here, we consider the simple model where a sequence S (e.g. a genome or a read) undergoes a simple mutation process whereby each nucleotide is mutated independently with some probability r, under the assumption that there are no spurious k-mer matches. How does this process affect the k-mers of S? We derive the expectation and variance of the number of mutated k-mers and of the number of islands (a maximal interval of mutated k-mers) and oceans (a maximal interval of non-mutated k-mers). We then derive hypothesis tests and confidence intervals for r given an observed number of mutated k-mers, or, alternatively, given the Jaccard similarity (with or without minhash). We demonstrate the usefulness of our results using a few select applications: obtaining a confidence interval to supplement the Mash distance point estimate, filtering out reads during alignment by Minimap2, and rating long read alignments to a de Bruijn graph by Jabba.

## 1 Introduction

*K*-mer-based methods have become widely used, e.g. for genome assembly [1], error correction [26], read mapping [16, 14], variant calling [31], genotyping [32, 7], database search [29, 11], metagenomic sequence comparison [37], and alignment-free sequence comparison [30, 22, 27]. A simple but influential recent example has been the Mash distance [22], which uses the minhash Jaccard similarity between the sets of *k*-mers in two sequences to estimate their average nucleotide divergence. Mash has been applied to determine the appropriate reference genome for in silico analyses [28], for genome compression [33], for clustering genomes [22, 3], and for estimating evolutionary distance from low-coverage sequencing datasets [27]. *K*-mer-based methods such as Mash are often faster and more practical then alignment-based methods. However, while the statistics behind sequence alignment are well understood [10], there are many gaps in our understanding of the statistics behind *k*-mer-based methods.

Consider the following simple mutation model and the questions it raises. There is a sequence of nucleotides *S* that undergoes a mutation process, whereby every position is mutated with some constant probability *r*_1_, independently of other nucleotides. In this model, we assume that *S* does not have any repetitive *k*-mers and that a mutation always results in a unique *k*-mer (we say that there are no *spurious matches*). This mutation model captures both a simple model of sequence evolution (e.g. Jukes-Cantor) and a simple model of errors generated during sequencing, under the assumptions that *k* is large enough and the repeat content low enough to make the effect of spurious matches negligible. It is applied to analyze algorithms and the predictions of the model often reflect performance on real biological sequences (e.g. [27, 22]).

How does this simple mutation model affect the *k*-mers of *S*? This question bears resemblance but is distinct from questions studied by Lander and Waterman [15] and in alignment-free sequence comparison [30] (we elaborate on the connection in Section 1.1). Some aspects of this question have been previously explored (e.g. [19, 26, 34]), but some very basic ones have not. For example, what is the distribution of the number of mutated *k*-mers? The expectation of this distribution is known and trivial to derive, but we do not know its variance. For another example, consider that the *k*-mers of *S* fall into mutated stretches (which, inspired by Lander-Waterman statistics, we call islands) and non-mutated stretches (which we call oceans). What is the distribution on the number of these stretches? We do not even know the expected value. We answer these and other questions in this paper, with most of the results captured in Table 1.

**Table 1:** The expectation, variances, and hypothesis tests derived in this paper. We use *q* as shorthand for 1*−*(1*−r*_1_)^*k*^. We use *f* (*r*_1_, *k*) as a placeholder for some function of *r*_1_ and *k* that is independent of *L*; see the theorems for the full expressions.

| Variable | Expectation | Variance | $(1 - \alpha)$ interval |
| --- | --- | --- | --- |
| $N_{\text{mut}}$ | $Lq$ | $L(1 - q)(q(2k + \frac{2}{r_1} - 1) - 2k) + f(r_1, k)$ | $Lq \pm z_\alpha \sqrt{\text{Var}(N_{\text{mut}})}$ |
| $N_{\text{isl}}$ | $Lr_1(1 - q) + f(r_1, k)$ | $Lr_1(1 - q)(1 - r_1(1 - q)(2k + 1)) + f(r_1, k)$ | $E[N_{\text{isl}}] \pm z_\alpha \sqrt{\text{Var}(N_{\text{isl}})}$ |
| $N_{\text{ocean}}$ | $Lr_1(1 - q) + f(r_1, k)$ | $Lr_1(1 - q)(1 - r_1(1 - q)(2k + 1)) + f(r_1, k)$ | $E[N_{\text{ocean}}] \pm z_\alpha \sqrt{\text{Var}(N_{\text{ocean}})}$ |
| Jaccard | — | — | $\left( \frac{L - Lq - z_\alpha \sqrt{\text{Var}(N_{\text{mut}})}}{L + Lq + z_\alpha \sqrt{\text{Var}(N_{\text{mut}})}}, \frac{L - Lq + z_\alpha \sqrt{\text{Var}(N_{\text{mut}})}}{L + Lq - z_\alpha \sqrt{\text{Var}(N_{\text{mut}})}} \right)$ |
| minhash Jaccard | — | — | see Theorem 6 |
| $C_{\text{ber}}$ | $\frac{L(1 - q)(1 + r_1(k - 1)) + f(r_1, k)}{L + k - 1}$ | see Theorem 11 | $E[C_{\text{ber}}] \pm z_\alpha \sqrt{\text{Var}(C_{\text{ber}})}$ |

We immediately apply our findings to derive hypothesis tests and confidence intervals for *r*_1_ from the number of observed mutated *k*-mers, the Jaccard similarity, and the Jaccard similarity under minhash. Previously, none were known, even though point estimates from these had been frequently used (e.g. Mash). In order to do this, we observe that our random variables are *m*-dependent [13], which, roughly speaking, means that the only dependencies involve *k*-mers nearby in the sequence. We apply a technique called Stein’s method [25] to approximate these as Normal variables and thereby obtain hypothesis tests and confidence intervals.

We demonstrate the usefulness of our results using a few select applications: obtaining a confidence interval to supplement the Mash distance point estimate [22], filtering out reads during alignment by Minimap2 [16], and rating long read alignments to a de Bruijn graph by Jabba [19]. These examples illustrate how the use of the simple mutation model and the techniques from our paper could have potentially improved several widely used tools. Our technique can also be applied to new questions as they arise. Our code for computing all the intervals in this paper is freely available at https://github.com/medvedevgroup/mutation-rate-intervals.

### 1.1 Related work

Here we give more background on how our paper relates to other previous work.

#### Lander-Waterman statistics

There is a natural analogy between the stretches of mutated *k*-mers and the intervals covered by random clones in the work of Lander and Waterman [15]. Each error can be viewed as a random clone with fixed length *k*, and thus the islands in our study correspond to “covered islands” in theirs. However, their focus was to determine how much redundancy was necessary to cover all (or most) of a genomic sequence, which would correspond to how many nucleotide mutations are needed so that most of the *k*-mers in the sequence are mutated. In particular, they expect average coverage of the sequence by clones to be greater than 1, while in our study we expect the corresponding value, ≈ *k*(1 − (1 − *r*)^*k*^, to be much less than 1. Thus, the approximations applied in [15] do not hold in our case.

#### Alignment-free sequence comparison

In alignment-free analysis, two sequences are compared by comparing their respective vectors of *k*-mer counts [30]. Two such vectors can be compared in numerous ways, e.g. through the the *D*_2_ similarity measure, which can be viewed as a generalization of the number of mutated *k*-mers we study in this paper. However, in alignment-free analysis, both the underlying model and the questions studied are somewhat different. In particular, alignment-free analysis usually works with much smaller values of *k*, e.g. *k <* 10 [38]. This means that most *k*-mers are present in a sequence, and *k*-mers will match between and within sequences even if they are in different locations and not evolutionarily related. Our model and questions assume that these spurious matches are background noise that can be ignored (which is justifiable for larger *k*), while they form a crucial component of alignment-free analysis. As a result, much of the work in measuring expectation and variance in metrics such as *D*_2_ is done with respect to the distribution of the original sequences, rather than after a mutation process [23, 5]. Even when the mutation processes have been studied, they have typically been very different from the ones we consider here (e.g. the “common motif model” [23]). Later works [20, 24] did consider the simple mutation model that we study here, though still with a small *k*. Sequence similarity has also been estimated using the average common substring length between two sequences [12]. This is similar to the distribution of oceans that we study in our paper, but the difference is that oceans are both left- and right-maximal, while the common substrings considered by [12] and others are only right-maximal.

## 2 Preliminaries

Let *L >* 0 be a positive integer. Let [*L*] to denote the interval of integers {0, …, *L* − 1}, which intuitively captures positions along a string. Let *k >* 0 be a positive integer. The *k-span* at position 0 ≤ *i < L* is denoted as *K*_*i*_ and is the range of integers [*i, i* + *k* − 1] (inclusive of the endpoints). Intuitively, a *k*-span captures the interval of a *k*-mer. We think of [*L* + *k* − 1] as representing an interval of length *L* + *k* − 1 that contains *L k*-spans. To simplify the statements of the theorems, we will in some places require that *L* ≥ *k* (or similar), i.e. that the interval is of length at least 2*k* − 1. We believe this covers most practical cases of interest, but, if necessary, the results can be rederived without this assumption.

We define the *simple mutation model* as a random process that takes as input two integers *k >* 0 and *L >* 0 and a real-valued *nucleotide error rate* 0 *< r*_1_ *<* 1. For every position in [*L* + *k* − 1], the process *mutates* it with probability *r*_1_. A mutation at position *i* is said to *mutate* the *k*-spans *K*_max(0,*i*−*k*+1)_, …, *K*_*i*_. We define *N*_mut_ as a random variable which is the number of mutated *k*-spans. As shorthand notation, we use *q* ≜:. 1 − (1 − *r*_1_)^*k*^ to denote the probability that a *k*-span is mutated. Figure 1 shows an example.

**Figure 1:**
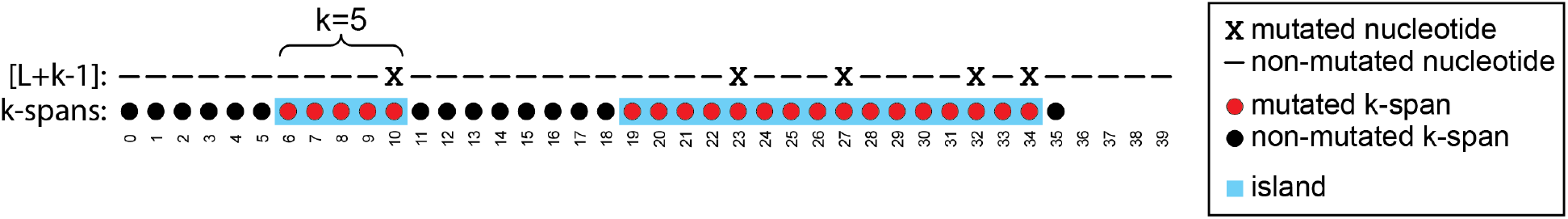
An example of the simple mutation process, with *L* = 36 and *k* = 5. There are 5 nucleotides that are mutated (marked with an x). For example, the mutation at position 10 mutates the *k*-spans *K*_6_, …, *K*_10_ (marked in red). Note that an isolated nucleotide mutation (e.g. at position 10) can affect up to *k k*-spans (e.g. *K*_6_, …, *K*_10_), but nearby nucleotide mutations can affect the same *k*-span (e.g. mutation of nucleotides at positions 23 and 27 both affect *K*_23_.) There are 2 islands (marked in blue) and 3 oceans, and *N*_mut_ = 21. For example, *K*_19_, …, *K*_34_ is an island, and *K*_35_ is an ocean.

The simple mutation model formalizes the notion of a string *S* undergoing mutations where there are no spurious matches, i.e. there are no duplicate *k*-mers in *S* and a mutation always creates a unique *k*-mer. This is also closely related to assuming that *S* is random and *k* is large enough so that such spurious matches happen with low probability. The simple mutation model captures these scenarios by representing *S* using the interval [*L* + *k* − 1] and a *k*-mer as a *k*-span.

We can partition the sequence *K*_0_, …, *K*_*L*−1_ into alternating intervals called *islands* and *oceans*. The range *i*, …, *j* is an *island* iff all *K*_*i*_, …, *K*_*j*_ are mutated, and the range is maximal, i.e. *K*_*i*−1_ and *K*_*j*+1_ are either not mutated or out of bounds. Similarly, the range is an *ocean* iff none of *K*_*i*_, …, *K*_*j*_ are mutated, and the interval is maximal. We define *N*_ocean_ as a random variable which is the number of oceans and *N*_isl_ as the number of islands (see Figure 1).

Consider two strings composed of a set of *k*-mers *A* and *B*, respectively, and let *s* ≤ min |*A*|, |*B*| be a non-negative integer. The *Jaccard similarity* between *A* and *B* is defined as 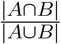. The *minhash sketch C*_*S*_ of a set *C* is the set of the *s* smallest elements in *C*, under a uniformly random permutation hash function. The *minhash Jaccard similarity* between *A* and *B* is defined as 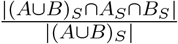, or, equivalently, |(*A* ∪ *B*)_*S*_ ∩ *A*_*S*_ ∩ *B*_*S*_|*/s* [2]. In order to transplant this to our model, we define the *sketching simple mutation model* as an extension of the simple mutation model, with an additional non-negative integer parameter *s* ≤ *L*. We follow the intuition of [*L* + *k* − 1] representing a string *S* with no spurious matches. For every position *i*, if *K*_*i*_ is non-mutated (respectively, mutated), we think of *K*_*i*_ as being shared (respectively, distinct) between the strings before and after the mutation process. Formally, let U be a universe which contains an element *shared*_*i*_ for every non-mutated *K*_*i*_ and, for every mutated *K*_*i*_, contains two elements *a*-*distinct*_*i*_ and *b*-*distinct*_*i*_. Let *A* be the set of all *shared*_*i*_ and *a*-*distinct*_*i*_, and let *B* be the set of all *shared*_*i*_ and *b*-*distinct*_*i*_. The output of the sketching simple mutation model is the minhash Jaccard similarity between *A* and *B*, i.e. Ĵ = |(*A* ∪ *B*)_*S*_ ∩ *A*_*S*_ ∩ *B*_*S*_|*/s*. Note that the Jaccard similarity (without sketches) would, in our simple mutation model, be the ratio between the number of *shared*_*i*_ and the size of U, which is 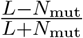.

Given a distribution with a parameter of interest *p*, an *approximate* (1 − *α*)*-confidence interval* is an interval which contains *p* with limiting probability 1 − *α*. Closely related, an *approximate hypothesis test with significance level* (1 − *α*) is an interval that contains a random variable with limiting probability 1 − *α*. We will drop the word “approximate” in the rest of the paper, for brevity. We will use the notation *X* ∈ *x* ± *y* to mean *X* ∈ [*x* − *y, x* + *y*]. Given 0 *< α <* 1, we define *z*_*α*_ = Φ^−1^(1 − *α/*2), where Φ^−1^ is the inverse of the cumulative distribution function of the standard Gaussian distribution. Let *H*(*x, y, z*) denote the hypergeometric distribution with population size *x, y* success states in population, and *z* trials. We define *F*_*n*_(*a*) = Pr[*H*(*L* + *n, L* −*n, s*) ≥*a*]. Both Φ^−1^ and *F*_*n*_ can be easily evaluated in programming languages such as R or python.

## 3 Number of mutated *k*-mers: expectation and variance

In this section, we look at the distribution of *N*_mut_, i.e. the number of mutated *k*-mers. The approach we take to this kind of analysis, which is standard, is to express *N*_mut_ as a sum of indicator random variables whose pairwise dependence can be derived. Let *X*_*i*_ be the 0*/*1 random variable corresponding to whether or not the *k*-span *K*_*i*_ is mutated; i.e., *X*_*i*_ = 1 iff at least one of its nucleotides is mutated. Hence, Pr[*X*_*i*_ = 1] = 1 − (1 − *r*_1_)^*k*^ ≜. *q*. We can express *N*_mut_ = Σ*X*_*i*_. By linearity of expectation, we have

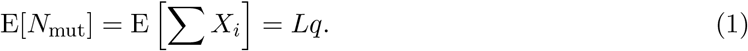

The key to the computation of variance is the joint probabilities of two *k*-mers being mutated.

### Lemma 1.

*Let* 0 ≤ *i < j < L. Then, X*_*i*_ *and X*_*j*_ *are independent if j* − *i* ≥ *k and* Pr[*X*_*i*_ = 1, *X*_*j*_ = 1] = 2*q* − 1 + (1 − *q*)(1 − *r*_1_)^*j*−*i*^ *otherwise*.

*Proof*. Set *δ* = *j* − *i*. If *δ* ≥ *k*, then *K_i_* and *K*_*i*+*δ*_ do not overlap and therefore the variables *X*_*i*_ and *X*_*i*+*δ*_ are independent. Otherwise, consider three events. *E*_1_ is the event that at least one of the positions *i*, …, *i* + *δ* − 1 is mutated. *E*_2_ is the event that none of *i*, …, *i* + *δ* − 1 is mutated and one of *i* + *δ*, …, *i* + *k* − 1 is mutated. *E*_3_ is the event that none of *i*, …, *i* + *k* − 1 is mutated. Notice that the three events form a partition of the event space and so we can write Pr[*X*_*i*_ = 1, *X*_*j*_ = 1] = Pr[*X*_*i*_ = 1, *X*_*j*_ = 1 | *E*_1_]Pr[*E*_1_] + Pr[*X*_*i*_ = 1, *X*_*j*_ = 1 | *E*_2_]Pr[*E*_2_] + Pr[*X*_*i*_ = 1, *X*_j_ = 1 | *E*_3_]Pr[*E*_3_] = Pr[*X*_j_ = 1 | *E*_1_]Pr[*E*_1_] + 1 · Pr[*E*_2_] + 0 · Pr[*E*_3_] = *q*(1 − (1 − *r*_1_)^*δ*^) + (1 − *r*_1_)^*δ*^(1 − (1 − *r*_1_)^*k*−*δ*^) = *q* − *q*(1 − *r*_1_)^*δ*^ + (1 − *r*_1_)^*δ*^ − (1 − *q*) = 2*q* − 1 + (1 − *q*)(1 − *r*_1_)^*δ*^.

We can now compute the variance using tedious but straightforward algebraic calculations. As we will show in the following section, knowing the variance allows us to obtain a confidence interval or do a hypothesis test based on *N*_mut_.

### Theorem 2.

*If* 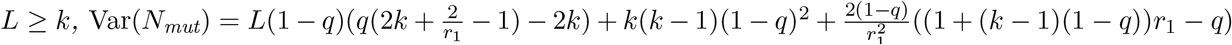.

## 4 Hypothesis test for *M* -dependent variables

Our derivations of hypothesis tests and confidence intervals follows the strategy used for the Binomials, which we now describe so as to provide intuition. In the case of estimating the success probability *p* of a Binomial variable *X* when the number of trials *L* is known, a confidence interval for *p* is called a binomial proportion confidence interval [6]. There are multiple ways to calculate such an interval, as described and compared in [4], and we will follow the approach of the Wilson score interval [36]. It works by first approximating the Binomial with a Normal distribution and then applying a standard score. The result is that 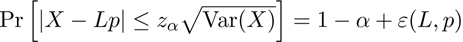, where Var(*X*) = *Lp*(1 − *p*) and *ε*(*L, p*) is a function such that lim_*L*→∞_ *ε*(*L, p*) = 0; recall that *z*_*α*_ = Φ^−1^(1−*α/*2). This can be solved for *X* to obtain a hypothesis test 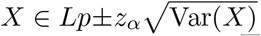. This can be converted into a confidence interval by finding all values of *p* for which 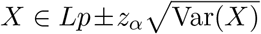 holds. In the particular case of the Binomial, a closed form solution is possible [36], but, more generally, one can also find the solution numerically.

Though random variables like *N*_mut_ are not Binomial, they have a specific form of dependence between the trials, which allows us to apply a similar strategy. A sequence of *L* random variables *X*_0_, …, *X*_*L*−1_ is said to be *m*-dependent if there exists a bounded *m* (with respect to *L*) such that if *j* − *i > m*, then the two sets {*X*_0_, …, *X*_*i*_} and {*X*_*j*_, …, *X*_*L*−1_} are independent [13]. In other words, *m*-dependence says that the dependence between a sequence of random variables is limited to be within blocks of length *m* along the sequence. It is known that the sum of *m*-dependent random variables is asymptotically normal [13] and this was previously used to construct heuristic hypothesis tests and confidence intervals [18]. Even stronger, the rate of convergence of the sum of *m*-dependent variables to the Normal distribution is known due to a technique called Stein’s method (see Theorem 3.5 in [25]). (This technique applies even to the case where *m* is not bounded, but that will not be the case in our paper.) Here, we apply Stein’s method to obtain a formally correct hypothesis test together with a rate of convergence for a sum of *m*-dependent (not necessarily identically distributed) Bernoulli variables.

### Lemma 3.

*Let* 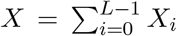 *be a sum of m-dependent Bernoulli random variables, where X*_*i*_ *has success probability p*_*i*_. *Let* 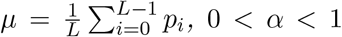, *and* 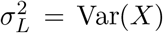. *Then*, Pr[*X* ≥ *Lµ* + *z*_*α*_*σ*_*L*_] = Pr[*X* ≤ *Lµ* − *z*_*α*_*σ*_*L*_] = *α/*2 − *ε/*2 *and*

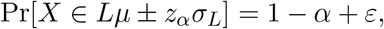

*where* 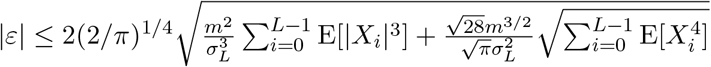.

*Proof*. Let *Y* = (*X* − *Lµ*)*/σ*_*L*_ and let *Z* be a standard normal random variable. From Theorem 3.6 in [25], we have 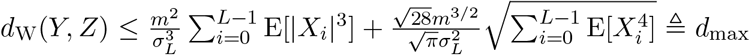, where *d*_W_(·, ·) denotes the Wasserstein metric. Since *Z* is a standard normal random variable, we have the following standard inequality between the Kolmogorov and Wasserstein metrics (see, e.g., Section 3 in [25]):

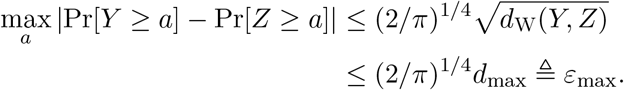

Recall that for a standard normal variable, Pr[*Z* ≥ *z*_*α*_] = *α/*2 and so, by the above, Pr[*Y* ≥ *z*_*α*_] ∈ *α/*2 ± *ε*_max_. Similarly, since Pr[*Z* ≤ −*z*_*α*_] = *α/*2 we obtain Pr[*Y* ≤ −*z*_*α*_] ∈ *α/*2 ± *ε*_max_. From the definition of *Y* it then follows that Pr[*X* ≥ *Lµ* + *z*_*α*_*σ*_*L*_] ∈ *α/*2 ± *ε*_max_ and Pr[*X* ≤ *Lµ* − *z*_*α*_*σ*_*L*_] ∈ *α/*2 ± *ε*_max_, and therefore implies that Pr[*X* ∈ *Lµ* ± *z*_*α*_*σ*_*L*_] ∈ 1 − *α* ± 2*ε*_max_.

As we will see, *m*-dependence is well-suited for dealing with variables in the simple mutation model. In most natural cases, the error |*ε*| → 0 when *L* → ∞, and Lemma 3 gives a hypothesis test with significance level 1 − *α*.

## 5 Hypothesis tests for *N*_mut_ and *Ĵ* and confidence intervals for *r*_1_

There is a natural point estimator for *r*_1_ using *N*_mut_, defined as 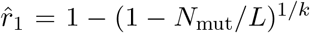. This estimator is both the method of moments and the maximum likelihood estimator, meaning it has nice convergence properties as *L* increases [35]. In this section, we extend it to a confidence interval and a hypothesis test, both from *N*_mut_ and *Ĵ* (with and without sketching). In the *N*_mut_ setting, Lemma 1 shows that *X*_0_, …, *X*_*L*−1_ are *m*-dependent with *m* = *k* − 1. Hence we can apply Lemma 3 to 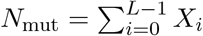.

### Corollary 4.

*Let* 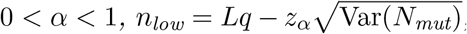, *and Let* 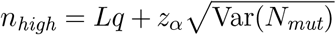. *Then* Pr[*N*_*mut*_ ≥ *n*_*high*_] = Pr[*N*_*mut*_ ≤ *n*_*low*_] = *α/*2 − *ε/*2 *and*

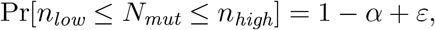

*where* |*ε*| ≤ *c/L*^1*/*4^ *and c is a constant that depends only on r*_1_ *and k. In particular, when r*_1_ *and k are independent of L, we have* lim_*L*→∞_(1 − *α* + *ε*) = 1 − *α*.

Corollary 4 gives the closed-form boundaries for a hypothesis test on *N*_mut_. To compute a confidence interval for *q* (equivalently, for *r*_1_), we can numerically find the range of *q* for which the observed *N*_mut_ lies between *n*_low_ and *n*_high_. In other words, the upper bound on the range would be given by the value of *q* for which the observed *N*_mut_ is *n*_low_ and the lower bound by the value of *q* for which the observed *N*_mut_ is *n*_high_. These observations are made rigorous in Theorem 5. We will use the notation 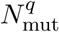 to denote *N*_mut_ with parameter *r*_1_ = 1 − (1 − *q*)^1*/k*^.

### Theorem 5.

*For fixed k, r*_1_, *and α, for a given observed value of* 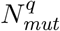, *there exists an L large enough such that there exists a unique q_low_ such that* 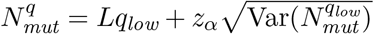 *and a unique q_high_ such that* 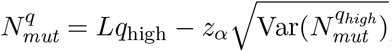, *and*

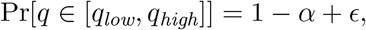

*where* |*ε*| ≤ *c/L*^1*/*4^ *and c is a constant that depends only on r*_1_ *and k. In particular, for fixed r*_1_ *and k, we have* lim_*L*→∞_(1 − *α* + *c*) = 1 − *α*.

Note that this theorem states that for sufficiently large *L*, there is a unique solution for the value of *q* for which the observed *N*_mut_ is *n*_high_ (and similarly a unique solution for the value of *q* for which the observed *N*_mut_ is *n*_low_). For small *L*, we have no such guarantee (though we believe the theorem holds true for all *L* ≥ *k*); to deal with this possibility, our software verifies if the solutions are indeed unique by computing the derivative inside the proof of Theorem 5 and checking if it is positive. If it is, then the proof guarantees the solutions to be unique; if it is not, our software reports this. However, during our validations, we did not find such a case to occur.

We want to underscore how the difference between a confidence interval and a hypothesis test is relevant in our case. A confidence interval is useful when we have two sequences, one of which having evolved from the other and we would like to estimate their mutation rate from the number of mutated *k*-spans. A hypothesis test is useful when we know the mutation rate a priori, e.g. the error rate of a sequencing machine. In this case, we may want to know whether a read could have been generated from a putative genome location, given the number of observed mutated *k*-spans. We will see both applications in Section 7.

In some cases, *N*_mut_ is not observed but instead we observe another random variable *T* = *f* (*N*_mut_), where *f* (*x*) is a monotone function. For example, if *f* (*x*) = (*L* − *x*)*/*(*L* + *x*), then *T* is the Jaccard similarity between the original and the mutated sequence (in our model). In this case, a hypothesis test with significance level *α* is to check if *T* lies between *f* (*n*_low_) and *f* (*n*_high_). In addition to the Jaccard, [17] describe 14 other variables that are a function of *N*_mut_, *L*, and *k*. These are: Anderberg, Antidice, Dice, Gower, Hamman, Hamming, Kulczynski, Matching, Ochiai, Phi, Russel, Sneath, Tanimoto and Yule. We can apply our hypothesis test to any of these variables, as long as they are monotone with respect to *N*_mut_.

We can also use Lemma 3 as a basis for deriving a hypothesis test on *Ĵ* in the sketching model. The proof is more involved and interesting in its own right, but is left for the Appendix due to space constraints.

### Theorem 6.

*Consider the sketching simple mutation model with known parameters s, k, L ≥ k, r*_1_, *and output JĴ. Let* 0 < *α* < 1 *and let m* ≥ 2 *be an integer*. *For* 0 ≤ *i* ≤ *m, let* 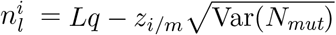 *and* 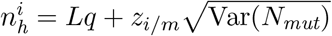. *Let*

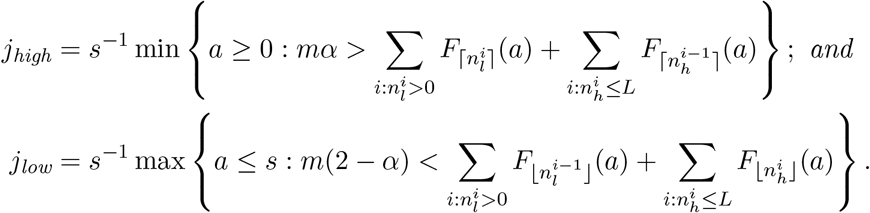

*Then, assuming that r*_1_ *and k are independent of L, and m* = *o*(*L*^1*/*4^),

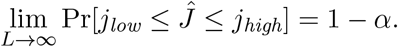

We can compute a confidence interval for *q* from *Ĵ* in the same manner as with Corollary 4. Let *j*_low_(*q*) and *j*_high_(*q*) be defined as in Theorem 6, but explicitly parameterized by the value of *q*. Then we numerically find the smallest value 0 *< q*_low_ *<* 1 for which *j*_low_(*q*_low_) = *Ĵ* and the largest value 0 *< q*_high_ *<* 1 for which *j*_high_(*q*_high_) = *Ĵ*. The following theorem guarantees that [*q*_low_, *q*_high_] is a confidence interval for *q*.

### Theorem 7.

*For fixed k, r*_1_, *α, m, and a given observed value of Ĵ, there exists an L large enough such that there exist unique intervals* 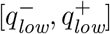 *and* 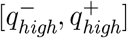 *such that* 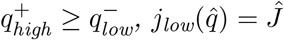 *if and only if* 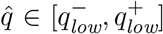, *and* 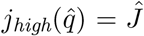 *if and only if* 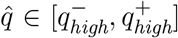. *Moreover, assuming that r*_1_, *k and m are independent of L, we have*

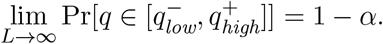

## 6 Number of islands and oceans

In this section, we derive the expectation and variance of *N*_isl_ and *N*_ocean_ and the hypothesis test based on them. For *N*_isl_, we follow the same strategy as for *N*_mut_, namely to express *N*_isl_ as a sum of indicator random variables whose joint probabilities can be derived. Let us define a *right border* as a position *i* such that *K*_*i*_ is mutated and *K*_*i*+1_ is not. We will denote it by an indicator variable *B*_*i*_, for 0 ≤ *i < L* − 1. Let us also say that there exists an *end-of-string border* iff *K*_*L*−1_ is mutated. We will denote this by an indicator variable *Z*. A right border is a position where an island ends and an ocean begins, and the end-of-string border exists if the last island is terminated not by an ocean but by the end of available nucleotides in the string to make a *k*-mer. The number of islands is then the number of borders, i.e. 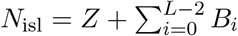.

To compute the expectation, observe that *Z* is a Bernoulli variable with parameter *q*. For *B*_*i*_, observe that the only way that *K*_*i*_ is mutated while *K*_*i*+1_ is not is if position *i* is mutated and the positions *i* + 1, …, *i* + *k* are not. Therefore, *B*_*i*_ ∼ *Bernoulli*(*r*_1_(1−*q*)). By linearity of expectation,

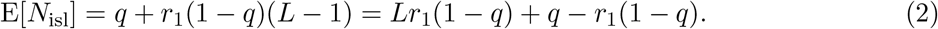

Next, we derive dependencies between border variables and use them to compute the variance.

### Lemma 8.

*Let* 0 ≤ *i < j* ≤ *L* − 2. *Then* Pr[*B*_*i*_ = 1, *B*_*j*_ = 1] = 0 *if j* ≤ *i* + *k and* 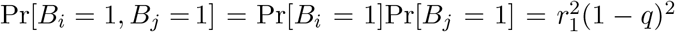 *otherwise. Also*, Pr[*B*_*i*_ = 1, *Z* = 1] = Pr[*B*_*i*_ = 1]Pr[*Z* = 1] = *r*_1_*q*(1 − *q*) *if i* ≤ *L* − 2 − *k, and* Pr[*B*_*i*_ = 1, *Z* = 1] = *r*_1_(1 − *q*)(1 − (1 − *r*_1_)^*L*−2−*i*^) *otherwise*.

*Proof*. Observe that when *j* − *i > k*, the positions that have an effect on *B*_*i*_ (i.e. *K*_*i*_, …, *K*_*i*+*k*_) and those that have an effect on *B*_*j*_ (i.e. *K*_*j*_, …, *K*_*j*+*k*_) are disjoint. Hence, *B*_*i*_ and *B*_*j*_ are independent in this case. When 1 ≤ *j* − *i* ≤ *k, B*_*i*_ and *B*_*j*_ cannot co-occur. This is because *B*_*i*_ = 1 implies that position *j* is not mutated, while *B*_*j*_ = 1 implies that it is. By the same logic, *Z* is independent of all *B*_*i*_ for 0 ≤ *i* ≤ *L* − 2 − *k*. For the case when *L* − 2 − *k < i* ≤ *L* − 2, *B*_*i*_ = 1 implies that positions *L* − 1, …, *i* + *k* are not mutated. Therefore, there is an end-of-string border when *B*_*i*_ = 1 iff one of the positions *i* + *k* + 1, …, *L* + *k* − 2 is mutated. Thus, Pr[*Z* = 1, *B*_*i*_ = 1] = Pr[*Z* = 1 | *B*_*i*_ = 1]Pr[*B*_*i*_ = 1] = (1 − (1 − *r*_1_)^*L*+*k*−2−(*i*+*k*+1)+1^)*r*_1_(1 − *q*).

### Theorem 9.

*For L* ≥ *k* +3, 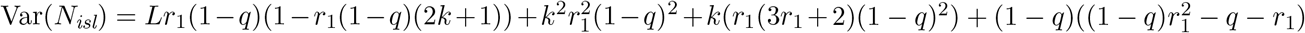.

Lemma 8 also shows that *N*_isl_ is *m*-dependent, with *m* = *k* − 1, Therefore, a hypothesis test on *N*_isl_ can be obtained as a corollary of Lemma 3.

### Corollary 10.

*Fix r*_1_ *and let* 0 *< α <* 1. *Then, the probability that* 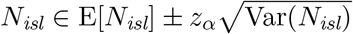 *is* 1 − *α* + *ε, where* |*ε*| ≤ *c/L*^1*/*4^ *and c is a constant that depends only on r*_1_ *and k. In particular, when r*_1_ *and k are independent of L, we have* lim_*L*→∞_(1 − *α* + *ε*) = 1 − *α*.

Unlike for Corollary 4, it is not as straightforward to invert this hypothesis test into a confidence interval for *r*_1_, since the endpoints of the interval of *N*_isl_ are not monotone in *r*_1_. We therefore do not pursue this direction here. The derivation of the expectation and variance for *N*_ocean_ is analogous and left for the Appendix (Theorem 12). Observe that |*N*_ocean_ − *N*_isl_| ≤ 1, so, as expected, the expectation and variance are identical to *N*_isl_ in the higher order terms. Corollary 10 also holds for the case that *n* is the observed number of oceans, if we just replace *N*_isl_ with *N*_ocean_.

An immediate application of *N*_ocean_ is to compute a hypothesis test for the *coverage by exact regions* (*C*_ber_), a variable earlier applied to score read mappings in [19]. *C*_ber_ is the fraction of positions in [*L* + *k* − 1] that lie in *k*-spans that are in oceans. The total number of bases in all the oceanic *k*-spans is the number of non-mutated *k*-spans plus, for each ocean, an extra *k* − 1 “starter” bases. We can then write

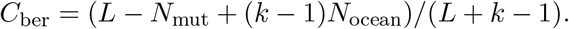

We can use the expectations and variances of *N*_mut_ (eq. (1) and Theorem 2) and *N*_ocean_ (Theorem 12) to derive the expectation and variance of *C*_ber_:

### Theorem 11.

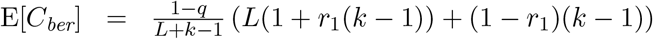*and, for L* ≥ *k* + 3,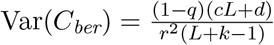, *where*

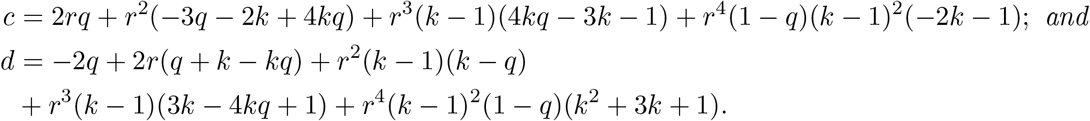

Then, observing that *C*_ber_ is a linear combination of *m*-dependent variables and hence itself *m*-dependent, we can apply Lemma 3 and obtain that, when *r*_1_ and *k* are independent of *L*,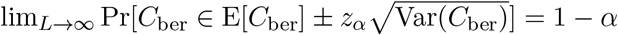.

## 7 Empirical results and applications

In this section, we evaluate the accuracy of our results and demonstrate several applications. A sanity check validation of the correctness of our formulas for E[*N*_mut_] and Var[*N*_mut_] is shown in Table S1, however, most of the expectation and variance formulas are evaluated indirectly through the accuracy of the corresponding confidence intervals. We focus the evaluation on accuracy rather than run time, since calculating the confidence interval took no more than a few seconds for most cases (the only exception was for sketch sizes of 100k or more, the evaluation took on the order of minutes). Memory use was negligible in all cases.

### 7.1 Confidence intervals based on *N*_mut_

In this section, we evaluate the accuracy of the confidence intervals (CIs) produced by Corollary 4 (other CIs will be evaluated indirectly through applications). We first simulate the simple mutation model to measure the accuracy, shown in the left three groups (i.e. *L* = 100, 1000, 10000) of Table 2, for *α* = 0.05. We observe that the predicted CIs are very accurate at *L* = 1000, and also accurate for smaller *k* and *r*_1_ when *L* = 100. Similar results hold for *α* = 0.01 (Table S2) and *α* = 0.10 (Table S3). The remainder of the cases had a CI that was too conservative; these are also the cases with some of the smallest variances (Table S1) and we suspect that, similar to the case of the Binomial, the Normal approximation of *m*-dependent variables deteriorates with very small variances. However, further investigation is needed.

**Table 2:** The accuracy of the confidence intervals for *r*_1_ predicted by Corollary 4, for *α* = 0.05 and for various values of *L, r*_1_, and *k* (the first three groups) and for the *E*.*coli* sequence (the fourth group). NA indicates the experiment was not run; for the first three groups, we only ran on parameters where ⌈E[*N*_mut_]⌉ < *L* (otherwise they were not of interest), while for *E*.*coli*, we ran with the same range of values of *r*_1_ and *k* as in the first three groups. In each cell, we report the fraction of 10,000 replicates for which the true *r*_1_ falls into the predicted confidence interval. For the *E*.*coli* sequence, we used strain Shigella flexneri Shi06HN159.

| | $L = 100$ | | | | $L = 1,000$ | | | | $L = 10,000$ | | | | *E. Coli* | | | |
| --- | --- | --- | --- | --- | --- | --- | --- | --- | --- | --- | --- | --- | --- | --- | --- | --- |
| $r_1 =$ | 0.001 | 0.01 | 0.1 | 0.2 | 0.001 | 0.01 | 0.1 | 0.2 | 0.001 | 0.01 | 0.1 | 0.2 | 0.001 | 0.01 | 0.1 | 0.2 |
| $k = 100$ | 0.91 | 1.00 | NA | NA | 0.95 | 0.96 | NA | NA | 0.95 | 0.95 | NA | NA | 0.95 | 0.95 | NA | NA |
| $k = 51$ | 0.91 | 1.00 | 1.00 | NA | 0.94 | 0.95 | 0.94 | NA | 0.95 | 0.95 | 0.96 | NA | 0.95 | 0.95 | 0.95 | NA |
| $k = 21$ | 0.91 | 0.96 | 1.00 | 1.00 | 0.93 | 0.95 | 0.95 | 0.95 | 0.95 | 0.94 | 0.95 | 0.95 | 0.95 | 0.94 | 0.93 | 0.94 |

Next, we investigate how well our predictions hold up when we simulate mutations along a real genome, where we can only observe the set of *k*-mers without their positions in the genome (as in alignment-free sequence comparison). We start with the *E*.*coli* genome sequence and, with probability *r*_1_, for every position, flip the nucleotide to one of three other nucleotides, chosen with equal probability. Let *A* and *B* be the set of distinct *k*-mers in *E*.*coli* before and after the mutation process, respectively. We let *L* = (|*A*| + |*B*|)*/*2 and *n* = *L* − |*A* ∩ *B*|. We then calculate the 95% CI for *r*_1_ under the simple mutation model (Corollary 4) by plugging in *n* for *N*_mut_. The rightmost group in Table 2 shows the accuracy of these CIs. We see that the simple mutation model we consider in this paper is a good approximation to mutations along a real genome like *E*.*coli*.

### 7.2 Mash distance

The Mash distance [22] (and its follow-up modifications [21, 27]) first measures the minhash Jaccard similarity *j* between two sequences and then uses a formula to give a point estimate for *r*_1_ under the assumptions of the sketching simple mutation model. While a hypothesis test was described in [22], it was only for the null model where the two sequences were unrelated. Theorem 6 allows us instead to give a CI for *r*_1_, based on the minhash Jaccard similarity, in the sketching simple mutation model. Table 3 reproduces a subset of Table 1 from [22], but using CIs given by Theorem 6. For most cases, the predicted CIs are highly accurate, with an error of at most two percentage points. The three exceptions happen when *s* is small and *q* is large; in such cases, the predicted CI is too conservative (i.e. too large). In Table S4, we also tested the accuracy with a real *E*.*coli* genome by letting *A* and *B* be the set of distinct *k*-mers in the genome before and after mutations, respectively, letting *L* = (|*A*| +|*B*|)*/*2 and *Ĵ* = ((|*A* ∪ *B*|) ∩ *A*_*s*_ ∩ *B*_*s*_)*/s*, and applying Theorem 6 with those values. The accuracy is very similar to that in the simple mutation model, demonstrating that for a genome like *E*.*Coli*, the simple mutation model is a good approximation.

**Table 3:**
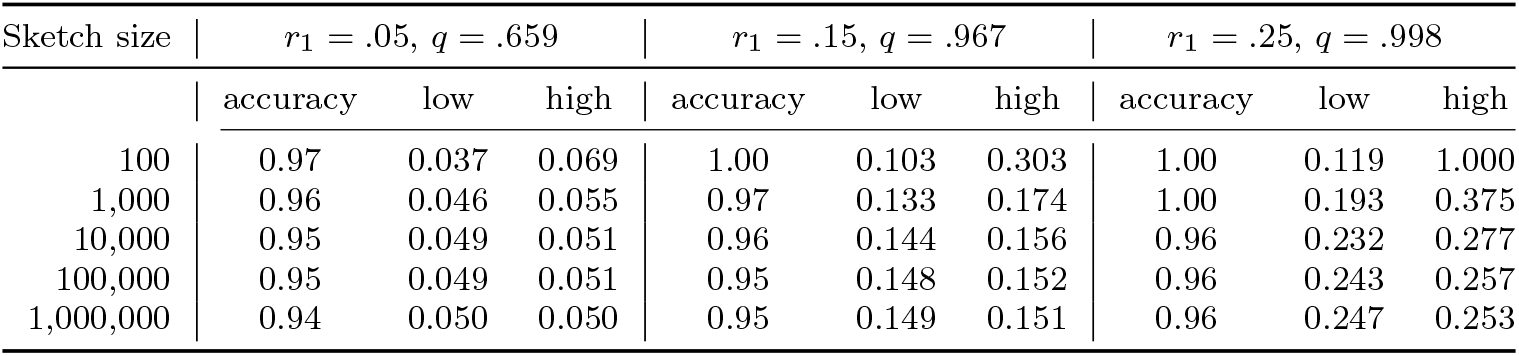
The confidence intervals predicted by Theorem 6 and their accuracy. For each sketch size and *r*_1_ value, we show the number of trials for which the true *r*_1_ falls within the predicted confidence interval. The reported CI corresponds to applying Theorem 6 with 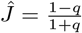. Here, *α* = 0.05, *k* = 21, *L* = 4, 500, 000, and the sketch size *s* and *r*_1_ are varied as shown. The number of trials for each cell is 1,000, and *m* = 100 for Theorem 6.

| Sketch size | $r_1 = .05, q = .659$ | | | $r_1 = .15, q = .967$ | | | $r_1 = .25, q = .998$ | | |
| --- | --- | --- | --- | --- | --- | --- | --- | --- | --- |
|  | accuracy | low | high | accuracy | low | high | accuracy | low | high |
| 100 | 0.97 | 0.037 | 0.069 | 1.00 | 0.103 | 0.303 | 1.00 | 0.119 | 1.000 |
| 1,000 | 0.96 | 0.046 | 0.055 | 0.97 | 0.133 | 0.174 | 1.00 | 0.193 | 0.375 |
| 10,000 | 0.95 | 0.049 | 0.051 | 0.96 | 0.144 | 0.156 | 0.96 | 0.232 | 0.277 |
| 100,000 | 0.95 | 0.049 | 0.051 | 0.95 | 0.148 | 0.152 | 0.96 | 0.243 | 0.257 |
| 1,000,000 | 0.94 | 0.050 | 0.050 | 0.95 | 0.149 | 0.151 | 0.96 | 0.247 | 0.253 |

### 7.3 Filtering out reads during alignment to a reference

Minimap2 is a widely used long read aligner [16]. The algorithm first picks certain *k*-mers in a read as *seeds*. Then, it identifies a region of the read and a region of the reference that potentially generated it (called a chain in [16]). Let *n* be the number of seeds in the read and let *m* ≤ *n* be the number of those that exist in the reference region. Minimap2 models the error rate of the *k*-mers as a homogenous Poisson process and estimates the sequence divergence between the read and the reference as 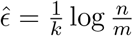 (which is the maximum likelihood estimator in that model). If 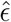 is above a threshold, the alignment is abandoned. [16] observes that due to invalid assumptions, 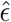 is only approximate and can be biased, but nevertheless maintains a good correlation with the true divergence.

Using our paper, we can obtain a more accurate estimate of *r*_1_. The situation is very similar to estimating *r*_1_ from *N*_mut_, except that only a subset of *k*-spans are being “tracked.” Therefore, the maximum likelihood estimator for *q* is *m/n* and for *r*_1_ is 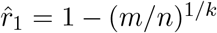. Figures 2 and S3 show the relative performance of the two estimators (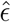 and 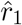) for sequences of different lengths, with our 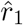 much closer to the simulated rate than 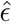 in both cases.

**Figure 2:**
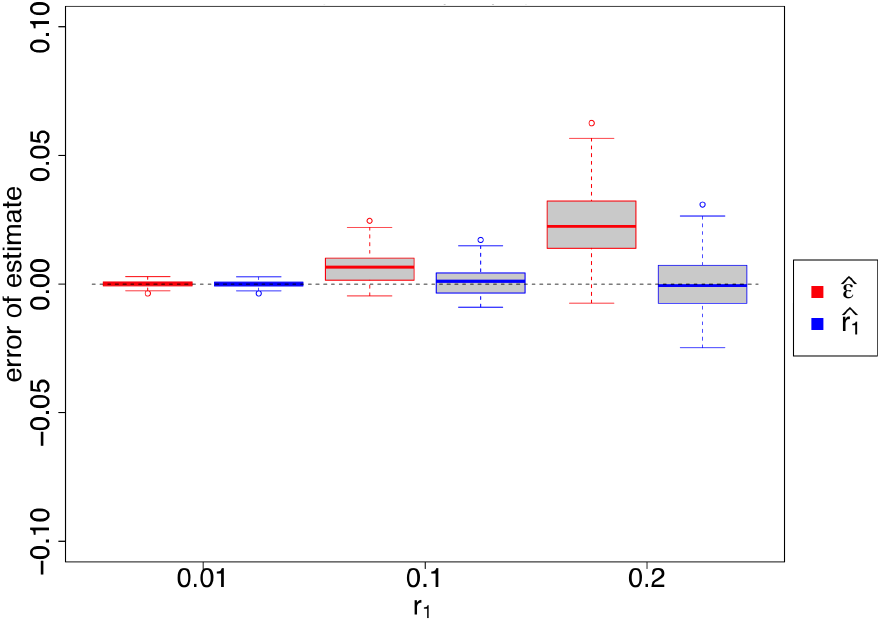
Estimates of sequence divergence as done by mimimap2 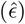 and by us 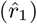. Reads are simulated from a random 10kbp sequence introducing mutations at the given *r*_1_ rate. For each *r*_1_ value, 100 reads are used. As in [16], we use *k* = 15 and, using a random hash function, identify as seeds the *k*-mer minimizers, one for every window of 25 *k*-mers. In the case when 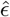 is undefined, we set 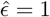.

### 7.4 Evaluating an alignment of a long read to a graph

Jabba [19] is an error-correction algorithm for long read data. At one stage, the algorithm evaluates whether a read is likely to have originated from a given location in the reference. Because Jabba’s reference is a de Bruijn graph and not a string, it uses the specialized *C*_ber_ score for the evaluation. In this scenario, the mutation process corresponds to sequencing errors at a known error rate *r*_1_ and the question is whether the read is likely to have arisen through this process from the given location of the reference. The authors assume the simple mutation model and derive the expected *C*_ber_ score as 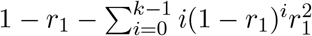. They then give a lower rating to reads with a *C*_ber_ score that has “significant deviation” from this expected value. It is not clear how much of a deviation is deemed to be significant or how it was calculated.

Theorem 11, which gives E[*C*_ber_] and Var[*C*_ber_], would have allowed [19] to take a more rigorous approach. It shows that the *C*_ber_ expectation computed by [19] is correct only in the limit as *L* → ∞, while our formula is exact and closed-form. More substantially, we can make the determination of “significant deviation” more rigorous. We regenerated Figure 2 from [19], using the same range of values for *k* (called *m* in [19]) and an error rate of *r*_1_ = 10% as in [19] and plotted the 95% confidence interval: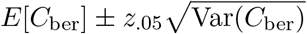. Figure 3 demonstrates that this range would have done a good job at capturing most of the generated reads. Table 4 gives the number of *C*_ber_ values that fall inside of the 95% confidence interval when using a simple mutation process with the same *r*_1_ = 10% for sequences of length 10,000 for 5,000 replicates, with *k* ranging from 5 to 50 in steps of 5, depicting good agreement between simulation and theorem 11.

**Figure 3:**
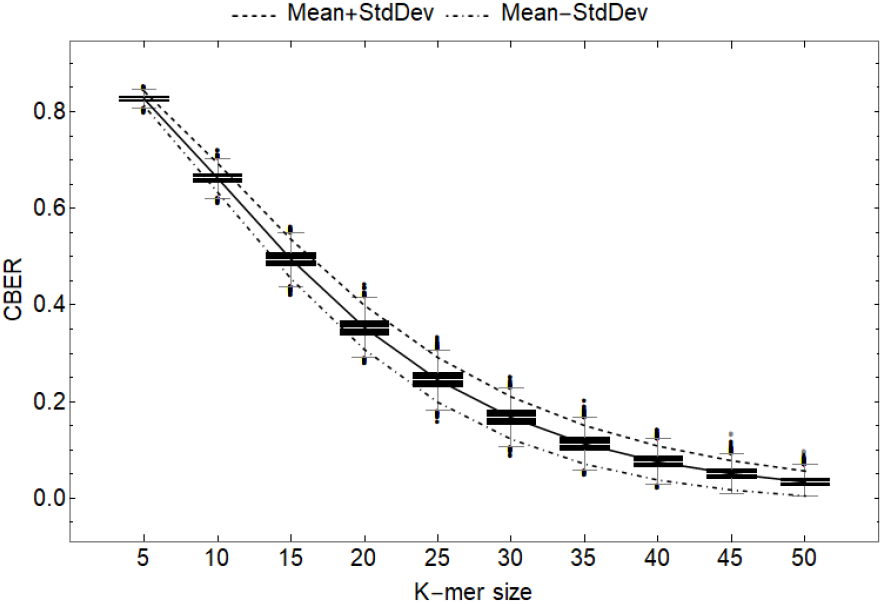
Box and whisker plot of *C*_ber_ scores for 5,000 replicates of random strings of length 10,000nt, with mutations introduced at a rate of *r*_1_ = 0.1. The solid black line corresponds to the empirical median of *C*_ber_, while the dashed top line corresponds to 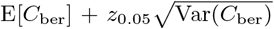 bottom dot-dashed line corresponds to and the 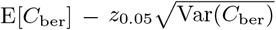, both computed from Theorem 11.

**Table 4:**
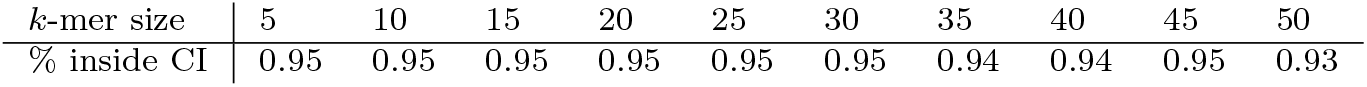
A total of 5,000 sequences, each of length 10,000nt underwent a simple mutation process with mutation probability *r*_1_ = 0.1. The percent of associated *C*_ber_ scores that fell inside of the 95% confidence interval as determined by Theorem 11 are shown.

| $k$-mer size | 5 | 10 | 15 | 20 | 25 | 30 | 35 | 40 | 45 | 50 |
| --- | --- | --- | --- | --- | --- | --- | --- | --- | --- | --- |
| % inside CI | 0.95 | 0.95 | 0.95 | 0.95 | 0.95 | 0.95 | 0.94 | 0.94 | 0.95 | 0.93 |

## 8 Conclusion

The simple mutation model has been used broadly to model either biological mutations or sequencing errors. However, its use has usually been limited to derive the expectations of random variables, e.g. the expected number of mutated *k*-mers. In this paper, we take this a step further and show that the dependencies between indicator variables in this model (e.g. whether a *k*-mer at a given position is mutated) are often easy to derive and are limited to nearby locations. This limited dependency allows us to show that the sum of these indicators is approximately Normal. As a result, we are able to obtain hypothesis tests and confidence tests in this model.

The most immediate application of our paper is likely to compute a confidence interval for average nucleotide identity from the minhash sketching Jaccard. Previously, only a point estimate was available, using Mash. However, we hope that our technique can be applied by others to random variables that we did not consider. All that is needed is to derive the joint probability of the indicator variables and compute the variance. Computing the variance by hand is tedious and error-prone but can be done with the aid of a software like Mathematica.

We test the robustness of the simple mutation model in the presence of spurious matches by using a real *E*.*coli* sequence. However, we do not explore the robustness with respect to violations such as the presence of indels (which result in different string lengths) or the presence of more repeats than in *E*.*coli*. This type of robustness has already been explored in other papers that use the simple mutation model [8, 22, 27]. However, exploring the robustness of our confidence intervals in downstream applications is important future work.

On a more technical note, it would be interesting to derive more tight error bounds for our confidence intervals, both in terms of more tightly capturing the dependencies on *L, r*_1_, and *k*, and accurately tracking constants. The error bound *ε* that is stated in Lemma 3 is likely not tight in either respect, due to the inherent loss when transferring between the Wasserstein and Kolmogorov metrics and due to loose inequalities within the proof of Theorem 3.5 in [25]. Ideally, tight error bounds would give the user a way to know, without simulations, when the confidence intervals are accurate, in the same way that we know that the Wilson score interval for a Binomial will be inaccurate when *np*(1 − *p*) is low. For example, it would be useful to better theoretically explain and predict which values in Table 2 deviate from 0.95.

Another practical issue is with the implementation of the algorithm to compute a confidence interval for *q* from Ĵ. Theorem 7 guarantees that the algorithm is correct as *L* goes to infinity. However, the user of the algorithm will not know if *L* is large enough for the confidence interval to be correct. There are several heuristic ways to check this, which we have implemented in the software: a short simulation to check the true coverage of the reported confidence interval, a check that the sets in the definitions of *j*_high_ and *j*_low_ are not empty, and a check that *j*_high_ and *j*_low_ are monotonic with respect to *q* in the range 0 *< q <* 1.

## Acknowledgements

PM is grateful to Kirsten E. Eilertson and Benjamin Shaby for discussion. PM was supported by NSF awards 1453527 and 1439057. AB was supported in part by NSF grant CCF-1850443. This material is based upon work supported by the National Science Foundation under Grant No. 1664803

## A Appendix

### A.1 Missing theorems and proofs

#### Theorem 2.

*If L* ≥ *k*, 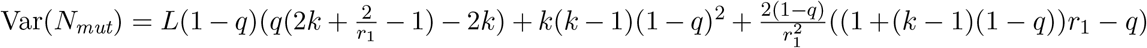.

*Proof*. In the following we will use Lemma 1 and the equality 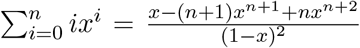 for *x* /= 1 from [9].

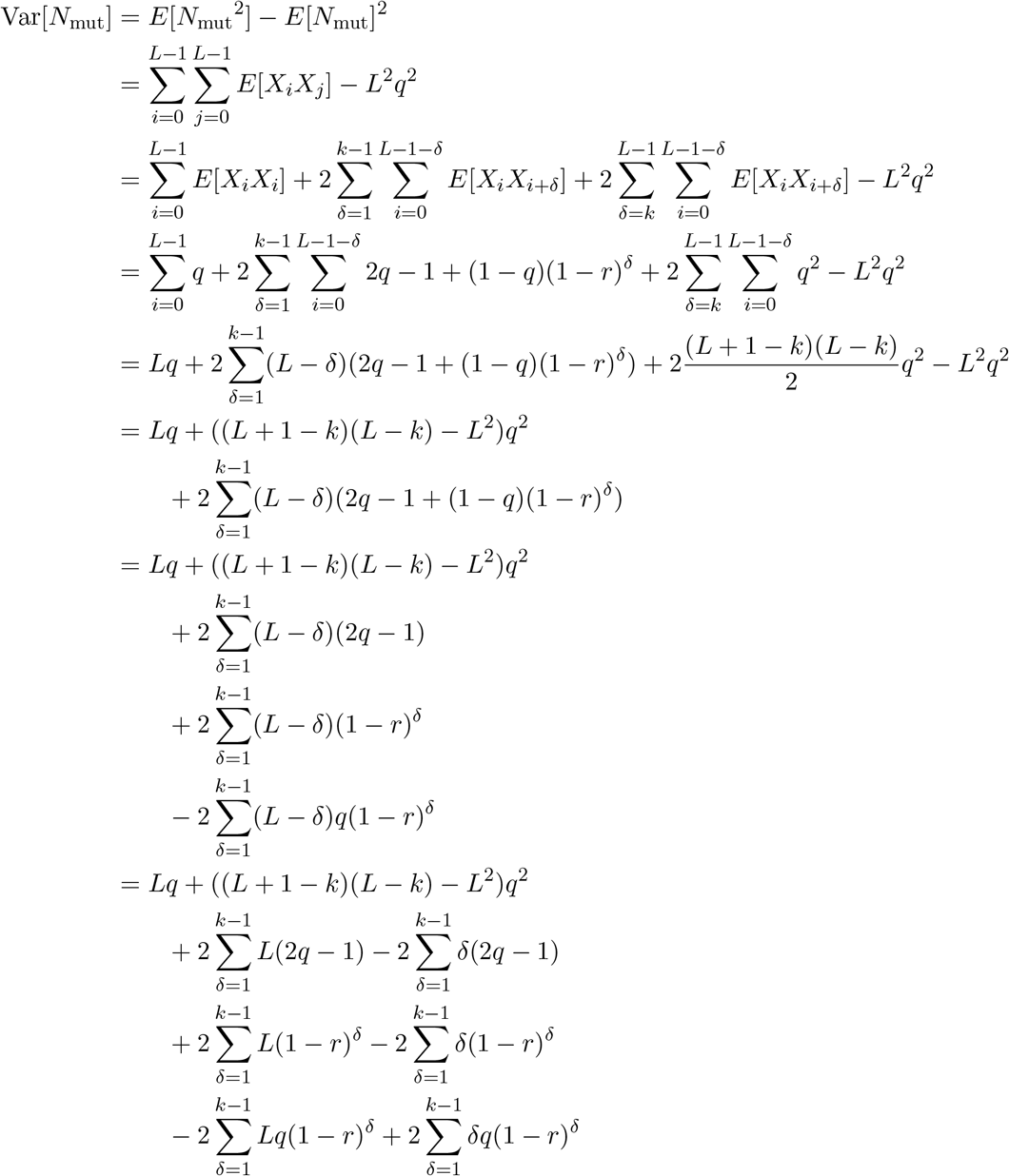

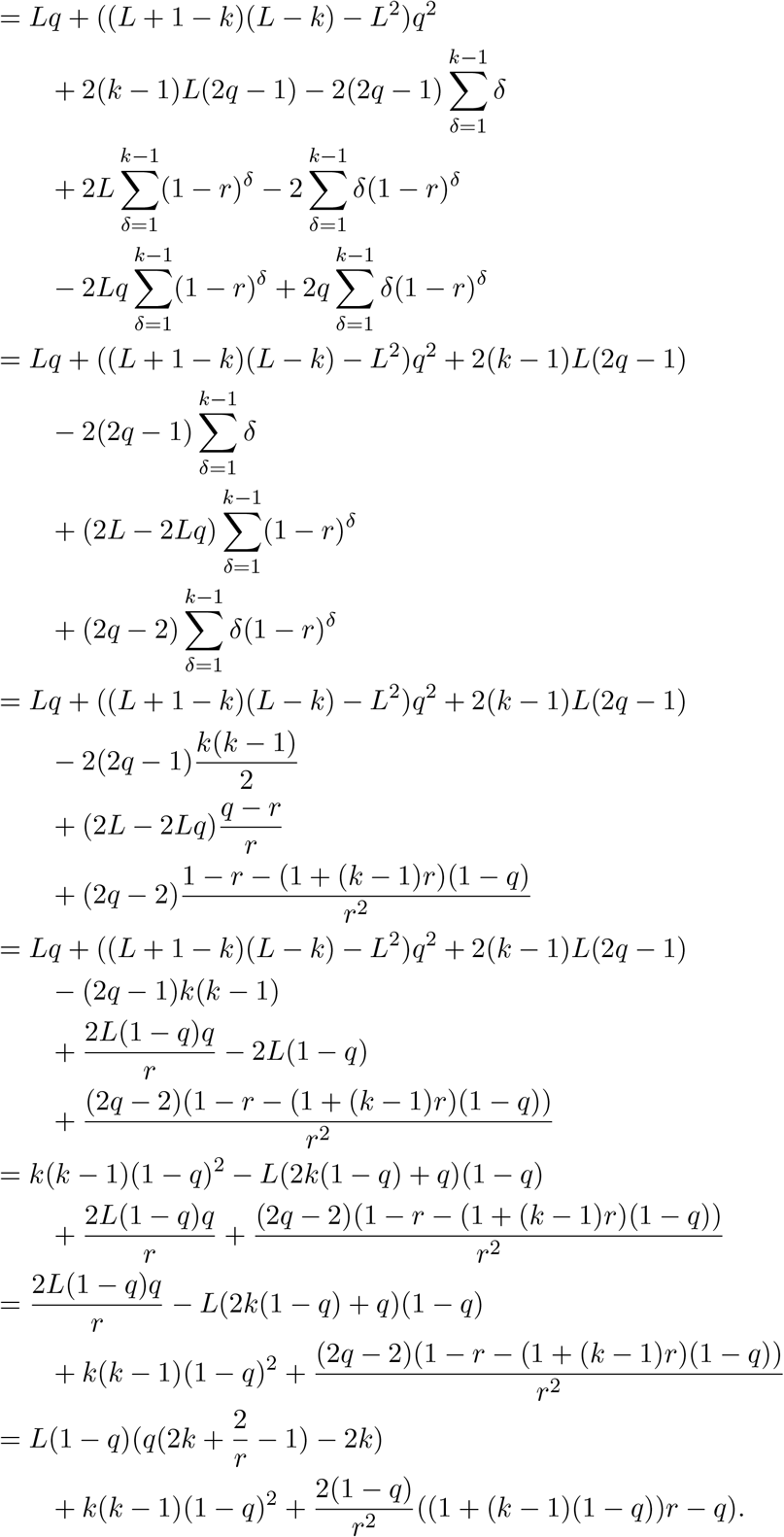

#### Theorem 5.

*For fixed k, r*_1_, *and α, for a given observed value of* 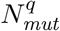, *there exists an L large enough such that there exists a unique q*_*low*_ *such that* 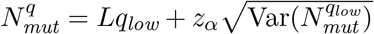 *and a unique q*_*high*_ *such that* 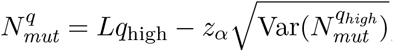, *and*

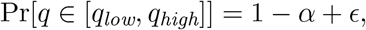

*Proof*. Given the result in corollary 4, we need only show that *q*_low_ and *q*_high_ are well-defined. As such, it is sufficient to show that 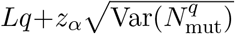 and 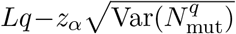 are strictly monotonic in *q* for sufficiently large *L*. Equivalently, since *q* = 1 − (1 − *r*_1_)^*k*^, these must be strictly monotonic in *r*_1_ which we consider here. For simplicity, we will write *r* instead of *r*_1_ and *N*_mut_ instead of 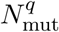. Focusing then on 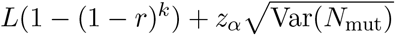, consider

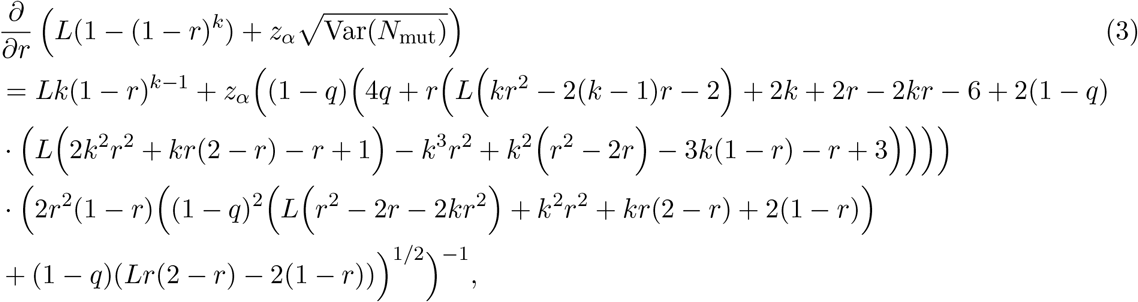

After a (tedious) series expansion of the rightside of the equality in eq. (3) about *L* = ∞, we find that 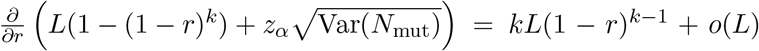. As such, 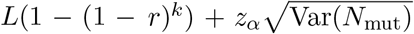 is increasing as a function of *r* as *L* → ∞. The case of showing that 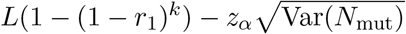 is also increasing proceeds in an entirely analogous fashion.

#### Theorem 6.

*Consider the sketching simple mutation model with known parameters s, k, L* ≥ *k, r*_1_, *and output* Ĵ. *Let* 0 *< α <* 1 *and let m* ≥ 2 *be an integer. For* 0 ≤ *i* ≤ *m, let* 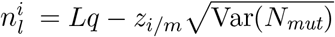 *and* 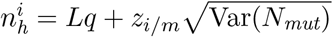. *Let*

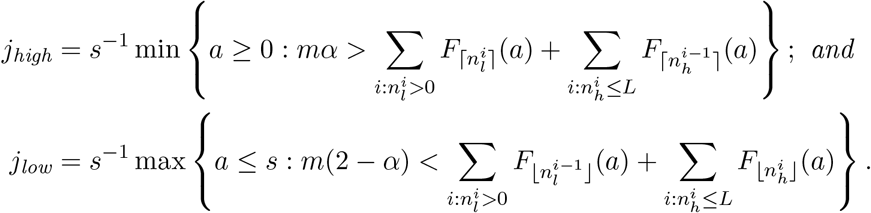

*Then, assuming that r*_1_ *and k are independent of L, and m* = *o*(*L*^1*/*4^),

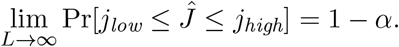

*Proof*. Recall the definition of *A* and *B* from the definition of the minhash Jaccard estimator. First, we argue that an element of (*A* ∪ *B*)_*S*_ is in *A*_*S*_ ∩ *B*_*S*_ iff it corresponds to a non-mutated *k*-span (i.e. iff *x* = *shared*_*i*_ for some *i*). Consider an element *x* ∈ (*A* ∪ *B*)_*S*_. If *x* corresponds to a mutated *k*-span (i.e. *x* = *a*-*distinct*_*i*_ or *x* = *b*-*distinct*_*i*_ for some *i*), then *x* ∉*A* ∩ *B* and so *x* ∉*A*_*S*_ ∩ *B*_*S*_. If *x* does not correspond to a mutated *k*-span (i.e. *x* = *shared*_*i*_ for some *i*), then *x* ∈ *A* ∩ *B* and *x* ∈ *A*_*S*_ ∩ *B*_*S*_ as well.

Next, let *J*′ be the random variable corresponding to |(*A* ∪ *B*)_*S*_ ∩ *A*_*S*_ ∩ *B*_*S*_|. The minhash Jaccard estimator can then be expressed as *ĵ* = *J′ /s*. Note that *J′* contains randomness due to both the mutation process and to the choice of the minhash permutation. For ease of notation we set *N* = *N*_mut_. We claim that the distribution of *J′*, conditioned on *N* = *n*, is hypergeometric *H*(*L* + *n, L* − *n, s*). To see this, recall from the discussion in the previous paragraphs that an element of (*A* ∪ *B*)_*S*_ is in *A*_*S*_ ∩ *B*_*S*_ only when it corresponds to a non-mutated *k*-span, and, on the event that *N* = *n*, there are exactly *L* − *n* non-mutated *k*-spans and a total of *L* + *n k*-spans to be hashed. We assume the hash values are assigned with a random permutation. Equivalently, we can generate the hash values by repeatedly assigning the smallest available hash value to an element chosen uniformly at random among those that have not been hashed yet. Then, from the first *s* elements, the probability that exactly *a* of those are selected from the set of the *L* − *n* non-mutated *k*-spans is:

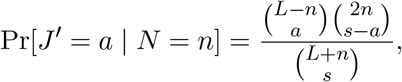

which corresponds to the hypergeometric *H*(*L* + *n*; *L*− *n, s*) probability function.

Our goal is to deduce a confidence interval for *J*Ĵ, or equivalently for *J*′. This can be easily done if we could compute:

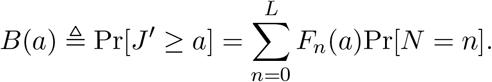

However, since we do not have an expression for Pr[*N* = *n*], we will instead obtain an upper and lower bound on *B*(*a*) using Lemma 3. For 1 ≤ *i* ≤ *m*, let 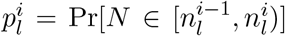 and let 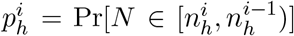. Note that 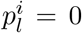 and 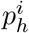 if 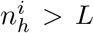. Using the law of total probability, we can write

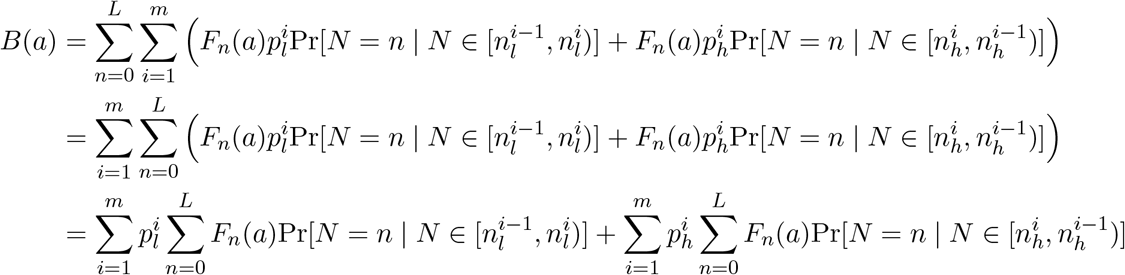

Observe that *F*_*n*_(*a*) is a non-increasing function with respect to *n*; this is because increasing *n* in *H*(*L* + *n*; *L* − *n, s*) has the overall effect of reducing the probability of success, since the population size is increased and the number of successes is decreased. Using this, we can find upper and lower bounds for *B*(*a*) as follows.

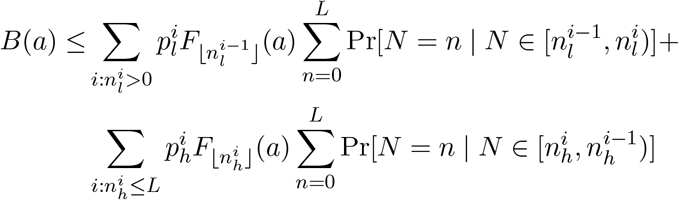

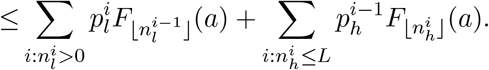

Similarly, we obtain 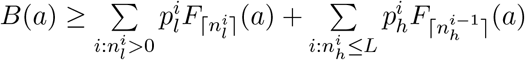.

We now approximate 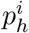 and 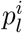 as follows. Observe that when 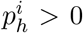, we have 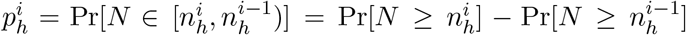. By Corollary 4, 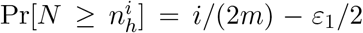 and 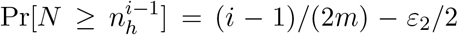; hence, 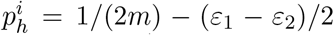. Here, *ε*_1_ and *ε*_2_ are constants whose absolute value is bounded by *ε*_max_ = *c/L*^1*/*4^, with *c >* 0 a constant that depends only on *q* and *k*. Hence, 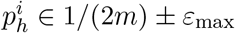 Analogously, we have 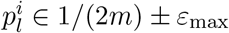.

This allows us to further simplify the bounds for *B*(*a*):

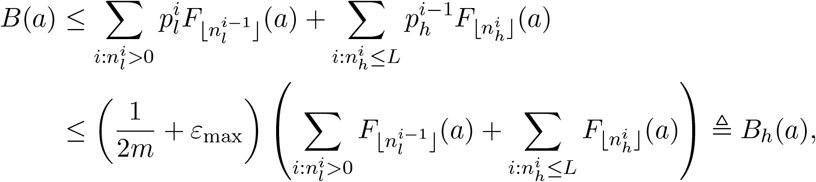

and

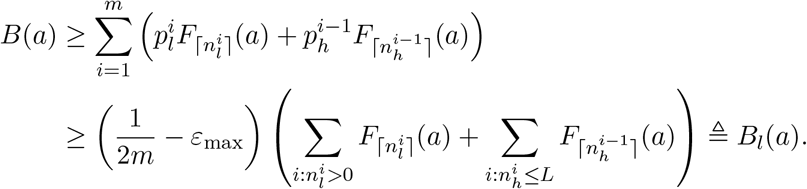

Let *a*_max_ = min{*a* ≥ 0: *α/*2 *> B*_*l*_(*a*)} and *a*_min_ = max{*a* ≤ *s*: *α/*2 *>* 1 − *B*_*h*_(*a*)}. Then, Pr[*J′* ∈ [*a*_min_, *a*_max_]] = 1 − *α* and so Pr[*J*Ĵ ∈ [*a*_min_*/s, a*_max_*/s*]] = 1 − *α*. The theorem then follows by observing that when *r*_1_, *k* are independent of *L* and *m* = *o*(*L*^1*/*4^), we have lim_*L*→∞_ *ε*_max_*m* = 0 and so *a*_min_*/s* → *j*_low_, and *a*_max_*/s* → *j*_high_.

#### Theorem 7.

*For fixed k, r*_1_, *α, m, and a given observed value of J*Ĵ, *there exists an L large enough such that there exist unique intervals* 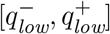 *and* 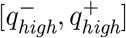 *such that* 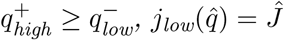 *if and only if* 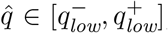, *and* 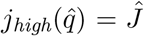 *if and only if* 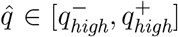. *Moreover, assuming that r*_1_, *k and m are independent of L, we have*

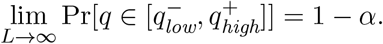

*Proof*. Recall from Theorem 6 that 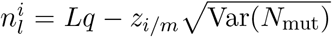 and 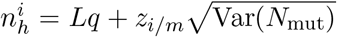. First, observe from Theorem 2 that Var(*N*_mut_) = *cL* + *o*(*L*), where *c* is a constant depending only on *k* and *r*_1_. Consequently, 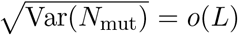 and so, for fixed *k, r*_1_, *α*, and *m*, there exists an *L* sufficiently large such that, for all 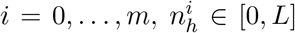 and 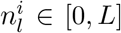. Therefore, for all values of *q*, there exists an *L* sufficiently large such that the summations in the definition of *j*_low_ and *j*_high_ are over 0 ≤ *i* ≤ *m*. Second, in the proof of Theorem 5 we established that 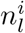 and 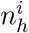 are increasing with *q* provided *L* is sufficiently large. Therefore, the parameters in the subscripts of the *F* terms of *j*_low_ and *j*_high_ are also increasing with *q*, when *L* is sufficiently large. Third, observe that *F*_*n*_(*a*) is a non-increasing function of *n* and of *a*. The fact that it is a non-increasing function of *n* we already observed in the proof of Theorem 6. The fact that it is a non-increasing function of *a* follows trivially from its definition. Combining these three observations, we deduce that *j*_low_ are *j*_high_ non-increasing functions of *q*. Therefore, they take on a certain value (i.e. *J*Ĵ) for a unique range of the domain, implying the first assertion of the theorem. The second assertion of the theorem then follows from Theorem 6.

**Figure S1:**
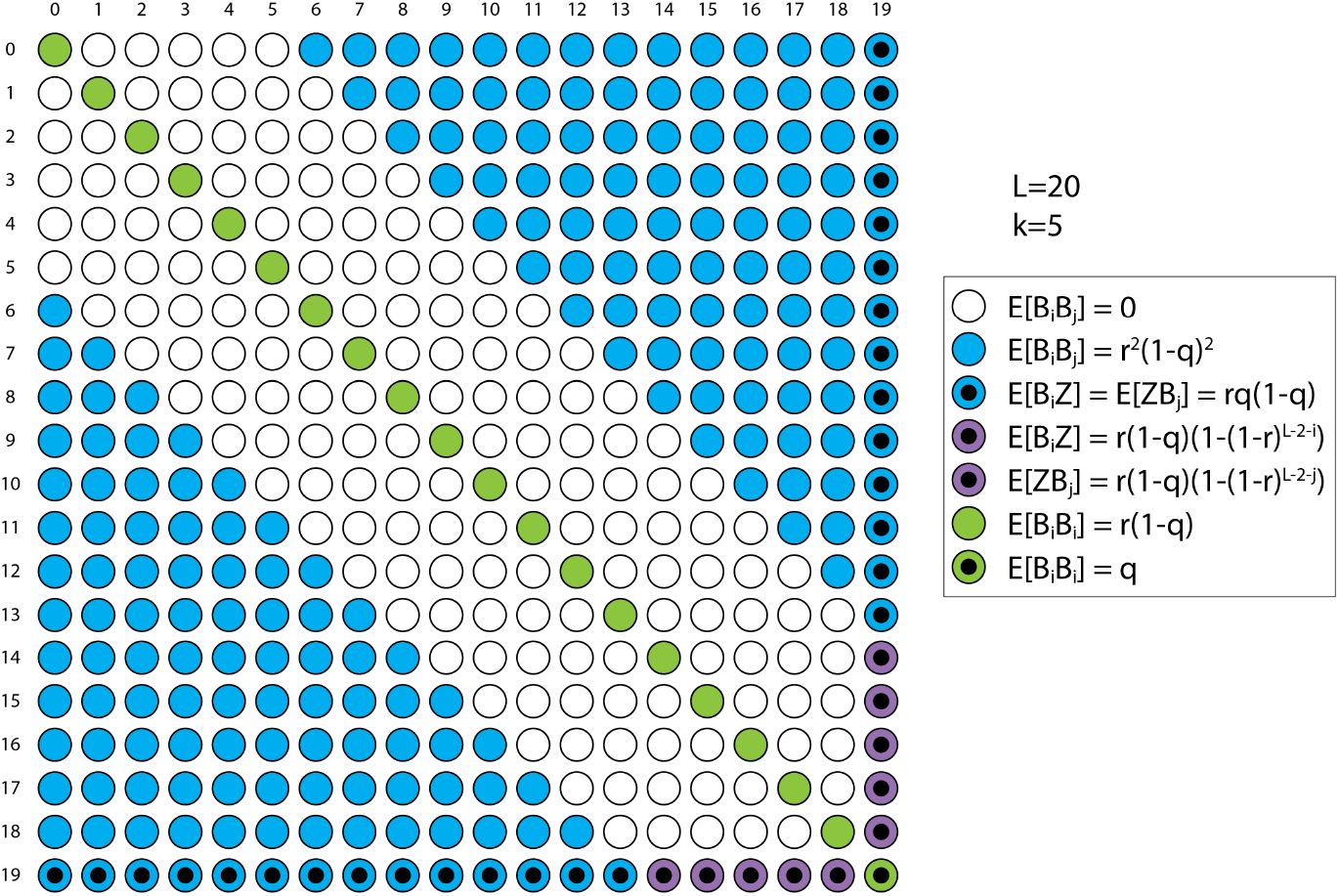
Illustration of the joint probabilities of *X*_*i*_ and *X*_*j*_, i.e. the terms of the sum in the derivation of Var(*N*_isl_) in Theorem 9. In this example, *L* = 20 and *k* = 5.

#### Theorem 9.

*For L* ≥ *k* +3,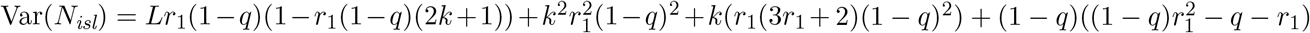.

*Proof*. For convenience, we will define a random variable *X*_*i*_ for 0 ≤ *i* ≤ *L* − 1 and let *X*_*i*_ = *B*_*i*_ for *i < L* − 1 and *X*_*L*−1_ = *Z*. Also, for notational simplicity, write *r* for *r*_1_. Figure S1 visualizes the joint probabilities of all *X*_*i*_’s, as given in Lemma 8. Using the figure as a guide, we proceed with the derivation.

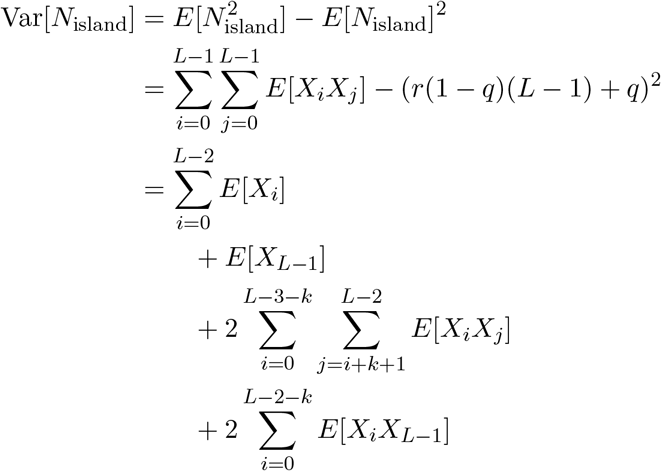

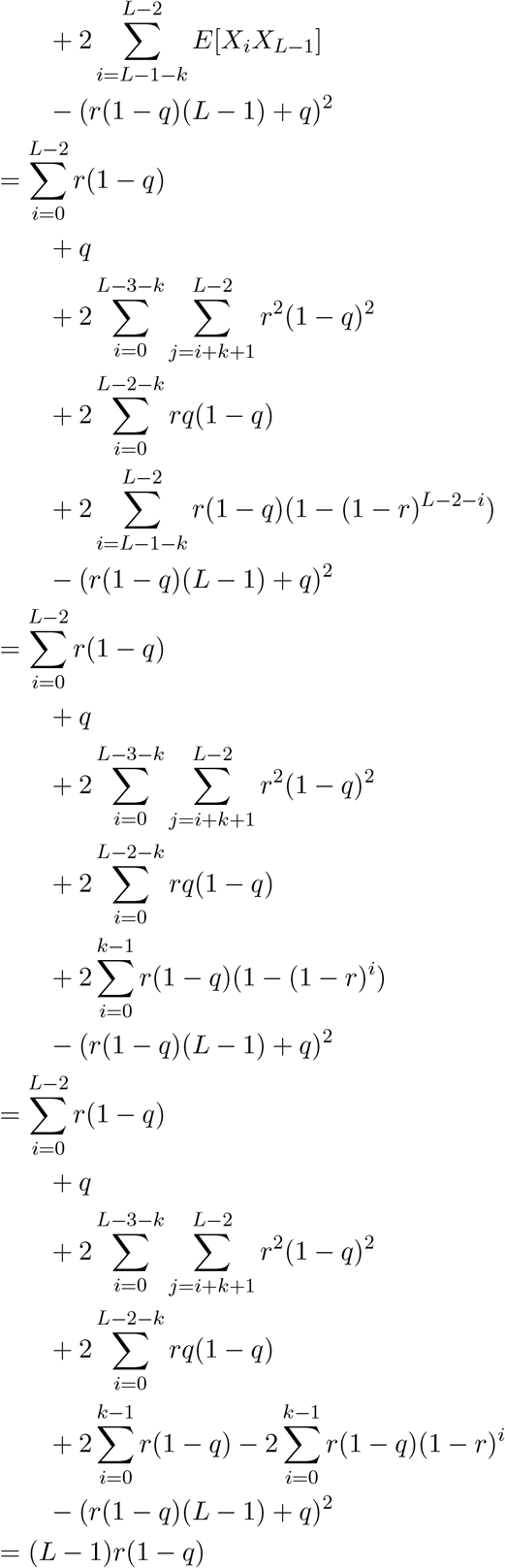

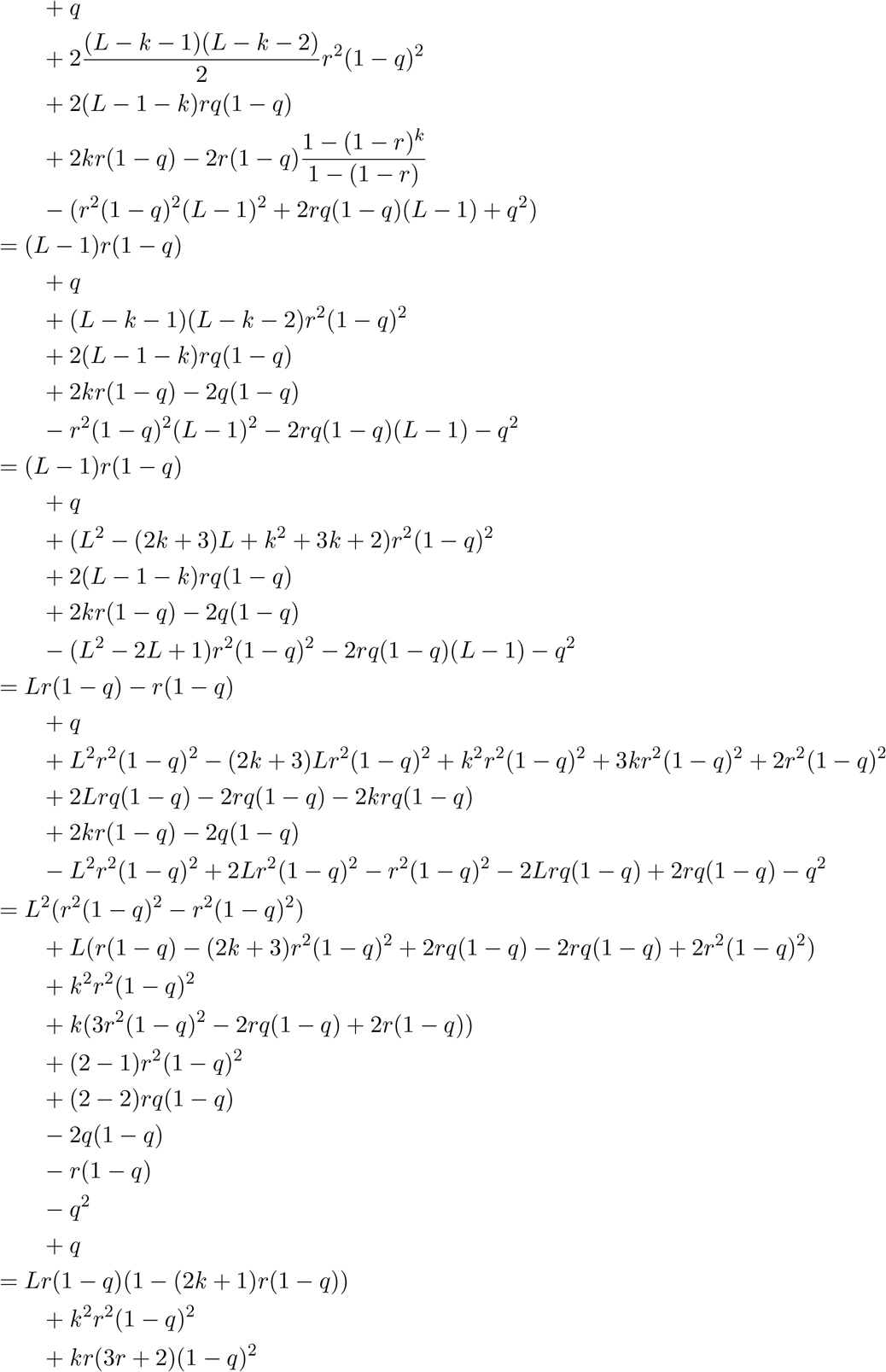

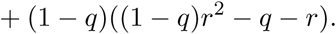

**Figure S2:**
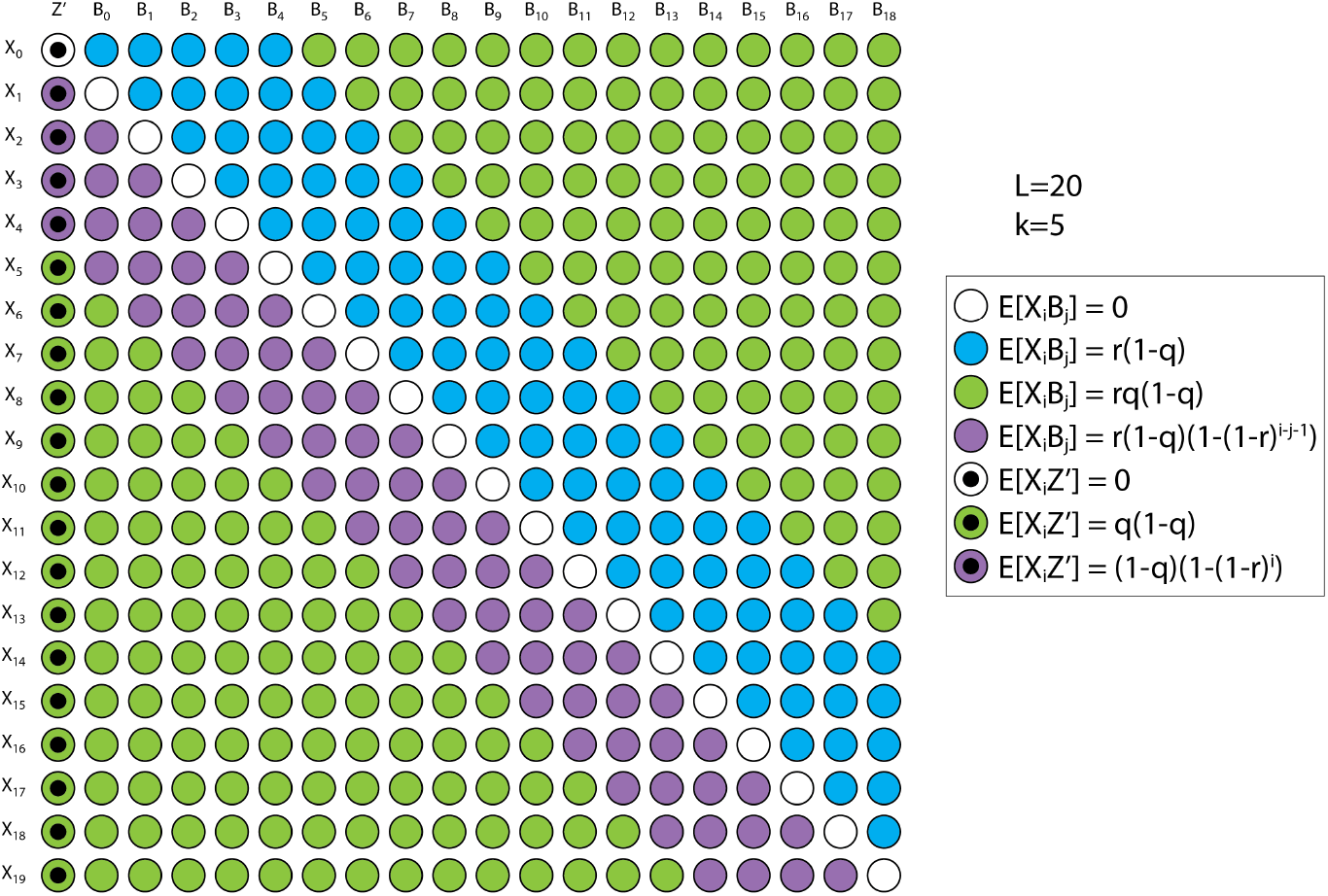
Illustration of the joint probabilities of *X*_*i*_, *B*_*j*_, and *Z′*, i.e. the terms of the sum in the derivation of Cov (*N*_ocean_, *N*_mut_) in Theorem 11. In this example, *L* = 20 and *k* = 5.

#### Theorem 12.

E[*N*_*ocean*_] = *Lr*_1_(1 − *q*) + (1 − *q*)(1 − *r*_1_) *and, for L* ≥ *k* + 3,

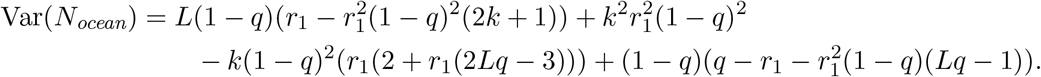

*Proof*. Let us define *Z′* as an indicator for the event that the first *k*-span (*K*_0_) is not mutated. Hence E[*Z′*] = (1 − *r*_1_)^*k*^ = (1 − *q*). Observe that every ocean begins either the start of the interval or a right border. Therefore, the the number of oceans is 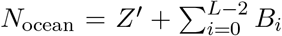. Thus E[*N*_ocean_] = (1 − *q*) +(*L* − 1)*r*_1_(1 − *q*) = (1 − *r*_1_)(1 − *q*) + *Lr*_1_(1 − *q*). For the variance, the derivation is equivalent to replacing *Z* with *Z* in the derivation of Theorem 9 and we therefore omit the proof here.

#### Theorem 11.

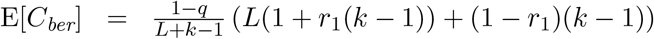*and, for L* ≥ *k* + 3, 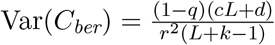, *where*

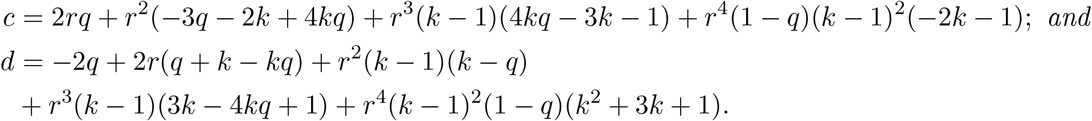

*Proof*. Throughout, we write *r* instead of *r*_1_ for simplicity. Recall that *C*_ber_ = (*L* − *N*_mut_ + (*k* − 1)*N*_ocean_)*/*(*L* + *k* − 1). Applying linearity of expectation together with eq. (1) and Theorem 12,

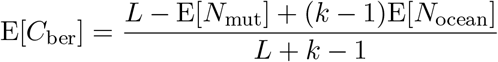

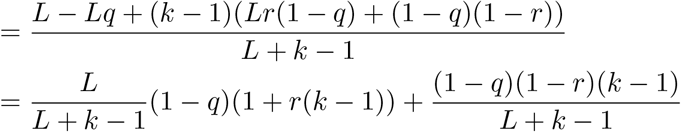

Applying the distributive properties of variance to the definition of *C*_ber_ we get:

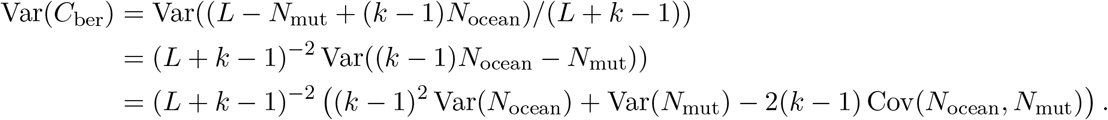

We only need to compute the covariance. We will use the same definition of variables as previously in Sections 3 and 6. In particular, *X*_*i*_ is a random variable indicating that *K*_*i*_ was mutated, *Z′* indicates that *K*_0_ has not mutated, and *B*_*i*_ indicates that *K*_*i*_ was mutated and *K*_*i*+1_ has not. Note that *Z′* = 1 − *X*_0_. Figure S2 visualizes the joint probabilities of all *X*_*i*_’s, *B*_*j*_’s, and *Z*. We then compute

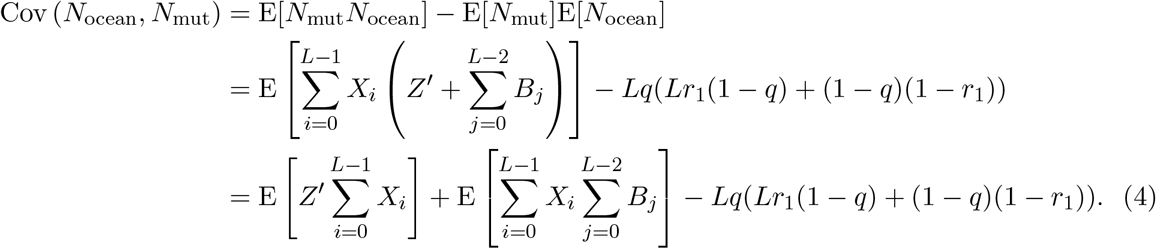

Observe that when *i* = 0, E[*Z′X*_*i*_] = 0. When 1 ≤ *i* ≤ *k* − 1, E[*Z*^t^*X*_*i*_] = (1 − *q*)(1 − (1 − *r*)^*i*^) since the left-most *i* nucleotides are not mutated when *Z′* = 1. When *k* ≤ *i* ≤ *L* − 1, *Z′* and *X*_*i*_ are independent so E[*Z′X*_*i*_] = *q*(1 − *q*). Thus, calculating the first sum in equation (4), we obtain

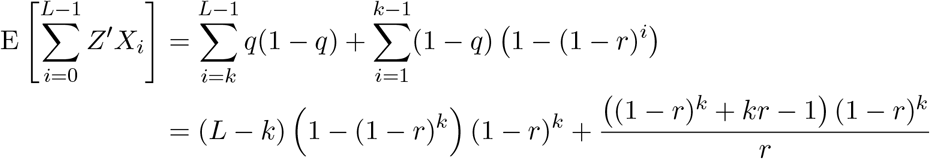

For the second sum in equation (4), observe that *B*_*j*_ = 1 implies that *X*_*j*_ = 1 and *X*_*j*+1_ = 0. Furthermore, for 2 ≤ *d* ≤ *k*, if *X*_*j*+*d*_ = 1, then the leftmost *k* + 1 + *d* nucleotides are unmutated, so from the law of total probability, E[*B*_*j*_*X*_*j*+*d*_] = (1 − (1 − *r*)^*d*−1^)*r*(1 − *q*). Lastly, if *B*_*j*_ = 1, then for all integers *i* such that max{0, *j* − *k* + 1} ≤ *i* ≤ *j, X*_*i*_ = 1 as well due to the mutation at position *j*. Hence E[*B*_*j*_*X*_*i*_] = E[*B*_*j*_] = *r*(1 − *q*). Using these observations, we obtain

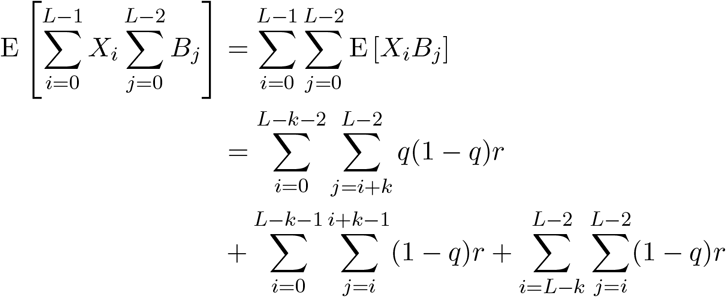

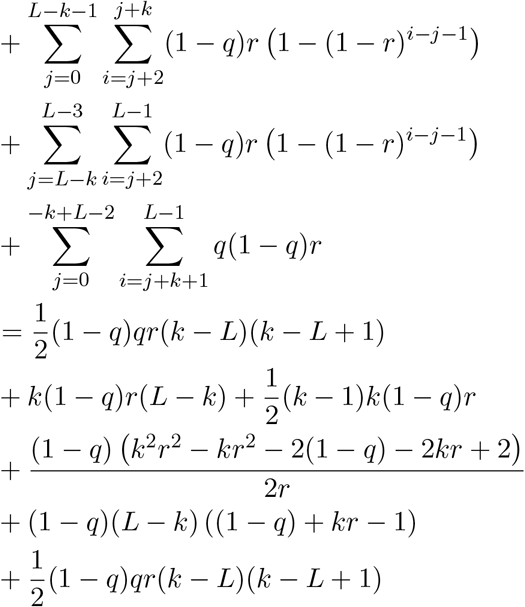

Putting all of this together and simplifying, we obtain

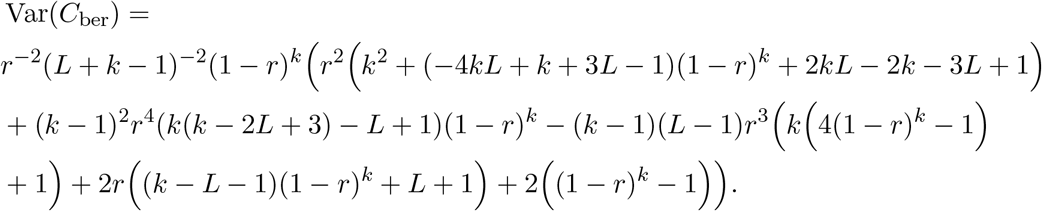

Factoring out the *L* terms in the numerator, we get

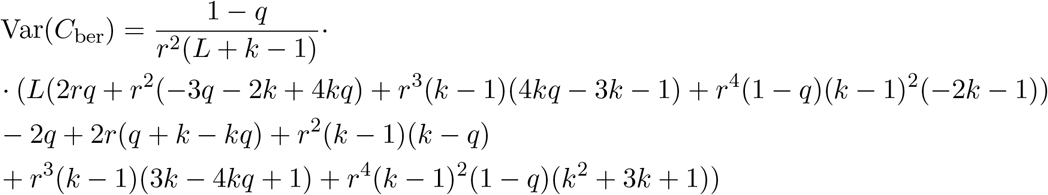

### A.2 Experimental results: extra tables and figures

**Table S1:**
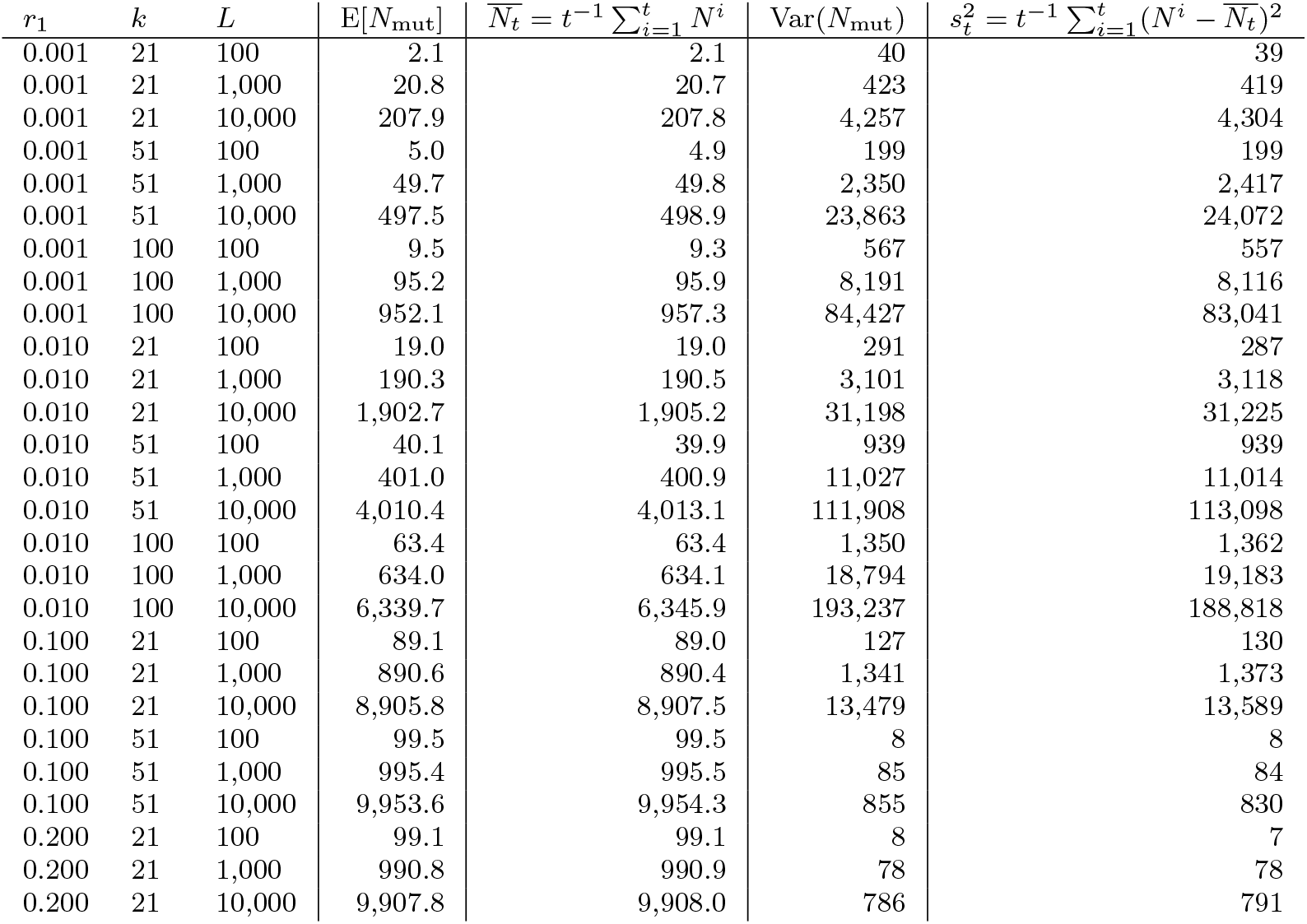
Validation of Equation (1) and Theorem 2, using *t* = 10, 000 trials. For each row, we show the value of E[*N*_mut_] given by Equation (1), the sample average of *N*_mut_ over all trials 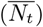, the value of Var[*N*_mut_] given by Theorem 2, and the sample variance of all the trials 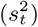. Here, *N* ^*i*^ is observed *N*_mut_ for the *i*^th^ trial.

**Table S2:**
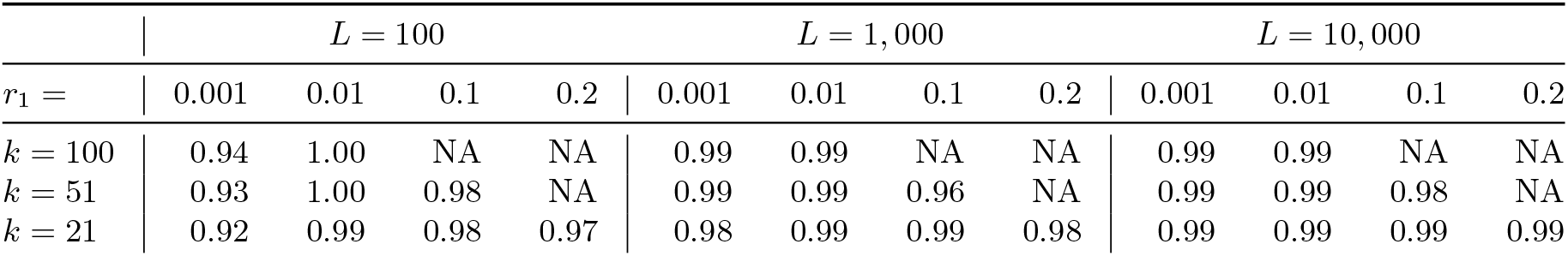
The accuracy of the confidence intervals for *r*_1_ predicted by Corollary 4, for *α* = 0.01 and for various values of *L, r*_1_, and *k*. NA indicates the experiment was not run because the parameters were not of interest (precisely, ⌈E[*N*_mut_]⌉ = *L*). The number of replicates was 10,000 for all experiments.

| | $L = 100$ | | | | $L = 1,000$ | | | | $L = 10,000$ | | | |
| --- | --- | --- | --- | --- | --- | --- | --- | --- | --- | --- | --- | --- |
| $r_1 =$ | 0.001 | 0.01 | 0.1 | 0.2 | 0.001 | 0.01 | 0.1 | 0.2 | 0.001 | 0.01 | 0.1 | 0.2 |
| $k = 100$ | 0.94 | 1.00 | NA | NA | 0.99 | 0.99 | NA | NA | 0.99 | 0.99 | NA | NA |
| $k = 51$ | 0.93 | 1.00 | 0.98 | NA | 0.99 | 0.99 | 0.96 | NA | 0.99 | 0.99 | 0.98 | NA |
| $k = 21$ | 0.92 | 0.99 | 0.98 | 0.97 | 0.98 | 0.99 | 0.99 | 0.98 | 0.99 | 0.99 | 0.99 | 0.99 |

**Table S3:** The accuracy of the confidence intervals for *r*_1_ predicted by Corollary 4, for *α* = 0.10 and for various values of *L, r*_1_, and *k*. NA indicates the experiment was not run because the parameters were not of interest (precisely, ⌈E[*N*_mut_]⌉ = *L*). The number of replicates was 10,000 for all experiments.

| | $L = 100$ | | | | $L = 1,000$ | | | | $L = 10,000$ | | | |
| --- | --- | --- | --- | --- | --- | --- | --- | --- | --- | --- | --- | --- |
| $r_1 =$ | 0.001 | 0.01 | 0.1 | 0.2 | 0.001 | 0.01 | 0.1 | 0.2 | 0.001 | 0.01 | 0.1 | 0.2 |
| $k = 100$ | 0.90 | 0.86 | NA | NA | 0.94 | 0.90 | NA | NA | 0.91 | 0.90 | NA | NA |
| $k = 51$ | 0.91 | 0.94 | 0.97 | NA | 0.93 | 0.90 | 0.93 | NA | 0.91 | 0.90 | 0.94 | NA |
| $k = 21$ | 0.91 | 0.94 | 0.92 | 0.94 | 0.93 | 0.90 | 0.90 | 0.93 | 0.92 | 0.90 | 0.90 | 0.90 |

**Table S4:**
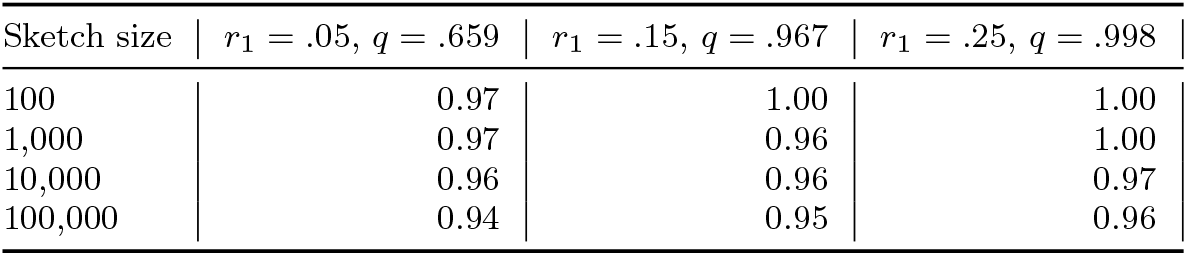
The accuracy of confidence intervals predicted by Theorem 6 on a real *E*.*coli* sequence. For each sketch size and *r*_1_ value, we show the number of trials for which the true *r*_1_ falls within the predicted confidence interval. Here, *α* = 0.05, *k* = 21, and the sketch size *s* and *r*_1_ are varied as shown. The number of trials for each cell is 1,000, and *m* = 100 for Theorem 6. *E*.*coli* strain K-12 substr. MG1655 was used.

**Figure S3:**
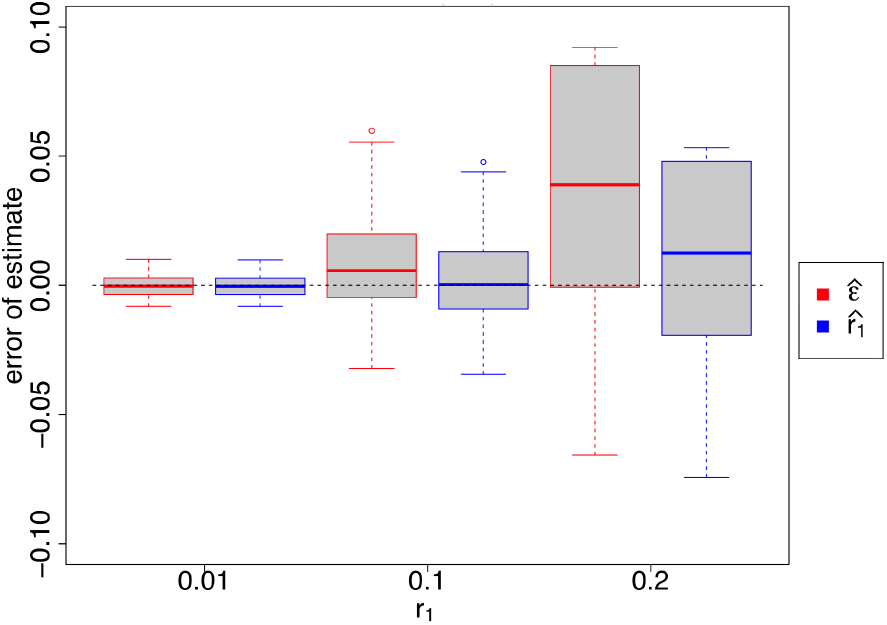
Estimates of sequence divergence as done by mimimap2 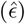 and by our approach 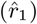. This is similar to Figure 2 but with sequence lengths of 1kbp instead of 10kbp.

## References

[1] Anton Bankevich, Sergey Nurk, Dmitry Antipov, Alexey A Gurevich, Mikhail Dvorkin, Alexander S Kulikov, Valery M Lesin, Sergey I Nikolenko, Son Pham, Andrey D Prjibelski, et al. SPAdes: a new genome assembly algorithm and its applications to single-cell sequencing. Journal of computational biology, 19(5):455–477, 2012.

[2] Andrei Z Broder. On the resemblance and containment of documents. In Proceedings. Compression and Complexity of SEQUENCES 1997 (Cat. No. 97TB100171), pages 21–29. IEEE, 1997.

[3] Christopher T Brown, Matthew R Olm, Brian C Thomas, and Jillian F Banfield. Measurement of bacterial replication rates in microbial communities. Nature biotechnology, 34(12):1256, 2016.

[4] Lawrence D Brown, T Tony Cai, and Anirban DasGupta. Interval estimation for a binomial proportion. Statistical science, pages 101–117, 2001.

[5] Conrad J Burden, Paul Leopardi, and Sylvain Forêt. The distribution of word matches between markovian sequences with periodic boundary conditions. Journal of Computational Biology, 21(1):41–63, 2014.

[6] George Casella and Roger L Berger. Statistical inference, volume 2. Duxbury Pacific Grove, CA, 2002.

[7] Luca Denti, Marco Previtali, Giulia Bernardini, Alexander Schönhuth, and Paola Bonizzoni. MALVA: genotyping by Mapping-free ALlele detection of known VAriants. iScience, 18:20–27, 2019.

[8] Huan Fan, Anthony R Ives, Yann Surget-Groba, and Charles H Cannon. An assembly and alignment-free method of phylogeny reconstruction from next-generation sequencing data. BMC genomics, 16(1):522, 2015.

[9] RL Grajam, Donald E Knuth, and Oren Patashnik. Concrete mathematics, a foundation for computer science, 1988.

[10] Dan Gusfield. Algorithms on strings, trees and sequences: computer science and computational biology. Cambridge University Press, 1997.

[11] R. S. Harris and P. Medvedev. Improved Representation of Sequence Bloom Trees. bioRxiv, 2018.

[12] Bernhard Haubold, Peter Pfaffelhuber, Mirjana Domazet-Loso, and Thomas Wiehe. Estimating mutation distances from unaligned genomes. Journal of Computational Biology, 16(10):1487–1500, 2009.

[13] Wassily Hoeffding, Herbert Robbins, et al. The central limit theorem for dependent random variables. Duke Mathematical Journal, 15(3):773–780, 1948.

[14] Chirag Jain, Alexander Dilthey, Sergey Koren, Srinivas Aluru, and Adam M Phillippy. A fast approximate algorithm for mapping long reads to large reference databases. In International Conference on Research in Computational Molecular Biology, pages 66–81. Springer, 2017.

[15] E. S. Lander and M. S. Waterman. Genomic mapping by fingerprinting random clones: a mathematical analysis. Genomics, 2(3):231–239, 1988.

[16] Heng Li. Minimap2: pairwise alignment for nucleotide sequences. Bioinformatics, 34(18):3094–3100, 2018.

[17] Yang Young Lu, Kujin Tang, Jie Ren, Jed A Fuhrman, Michael S Waterman, and Fengzhu Sun. Cafe: accelerated alignment-free sequence analysis. Nucleic acids research, 45(W1):W554–W559, 2017.

[18] Weiwen Miao and Joseph L Gastwirth. The effect of dependence on confidence intervals for a population proportion. The American Statistician, 58(2):124–130, 2004.

[19] Giles Miclotte, Mahdi Heydari, Piet Demeester, Stephane Rombauts, Yves Van de Peer, Pieter Audenaert, and Jan Fostier. Jabba: hybrid error correction for long sequencing reads. Algorithms for Molecular Biology, 11(1):1–12, 2016.

[20] Burkhard Morgenstern, Bingyao Zhu, Sebastian Horwege, and Chris André Leimeister. Estimating evolutionary distances between genomic sequences from spaced-word matches. Algorithms for Molecular Biology, 10(1):5, 2015.

[21] Brian D Ondov, Gabriel J Starrett, Anna Sappington, Aleksandra Kostic, Sergey Koren, Christopher B Buck, and Adam M Phillippy. Mash Screen: High-throughput sequence containment estimation for genome discovery. Genome biology, 20(1):232, 2019.

[22] Brian D Ondov, Todd J Treangen, Páll Melsted, Adam B Mallonee, Nicholas H Bergman, Sergey Koren, and Adam M Phillippy. Mash: fast genome and metagenome distance estimation using minhash. Genome biology, 17(1):132, 2016.

[23] Gesine Reinert, David Chew, Fengzhu Sun, and Michael S Waterman. Alignment-free sequence comparison (i): statistics and power. Journal of Computational Biology, 16(12):1615–1634, 2009.

[24] Sophie Röhling, Alexander Linne, Jendrik Schellhorn, Morteza Hosseini, Thomas Dencker, and Burkhard Morgenstern. The number of k-mer matches between two dna sequences as a function of k and applications to estimate phylogenetic distances. Plos one, 15(2):e0228070, 2020.

[25] Nathan Ross. Fundamentals of Stein’s method. Probability Surveys, 8:210–293, 2011.

[26] Leena Salmela, Riku Walve, Eric Rivals, and Esko Ukkonen. Accurate self-correction of errors in long reads using de Bruijn graphs. Bioinformatics, 33(6):799–806, 2017.

[27] Shahab Sarmashghi, Kristine Bohmann, M Thomas P Gilbert, Vineet Bafna, and Siavash Mirarab. Skmer: assembly-free and alignment-free sample identification using genome skims. Genome biology, 20(1):1–20, 2019.

[28] Oliver Schwengers, Torsten Hain, Trinad Chakraborty, and Alexander Goesmann. Reference-seeker: rapid determination of appropriate reference genomes. BioRxiv, page 863621, 2019.

[29] B. Solomon and C. Kingsford. Fast search of thousands of short-read sequencing experiments. Nature biotechnology, 34(3):300–302, 2016.

[30] Kai Song, Jie Ren, Gesine Reinert, Minghua Deng, Michael S Waterman, and Fengzhu Sun. New developments of alignment-free sequence comparison: measures, statistics and next-generation sequencing. Briefings in bioinformatics, 15(3):343–353, 2014.

[31] Daniel S Standage, C Titus Brown, and Fereydoun Hormozdiari. Kevlar: a mapping-free framework for accurate discovery of de novo variants. bioRxiv, page 549154, 2019.

[32] Chen Sun and Paul Medvedev. Toward fast and accurate snp genotyping from whole genome sequencing data for bedside diagnostics. Bioinformatics, 35(3):415–420, 2018.

[33] Tao Tang, Yuansheng Liu, Buzhong Zhang, Benyue Su, and Jinyan Li. Sketch distance-based clustering of chromosomes for large genome database compression. BMC genomics, 20(10):1–9, 2019.

[34] Anqi Wang and Kin Fai Au. Performance difference of graph-based and alignment-based hybrid error correction methods for error-prone long reads. Genome biology, 21(1):14, 2020.

[35] Larry Wasserman. All of statistics: a concise course in statistical inference. Springer Science & Business Media, 2013.

[36] Edwin B Wilson. Probable inference, the law of succession, and statistical inference. Journal of the American Statistical Association, 22(158):209–212, 1927.

[37] Derrick E Wood and Steven L Salzberg. Kraken: ultrafast metagenomic sequence classification using exact alignments. Genome biology, 15(3):R46, 2014.

[38] Tiee-Jian Wu, Ying-Hsueh Huang, and Lung-An Li. Optimal word sizes for dissimilarity measures and estimation of the degree of dissimilarity between dna sequences. Bioinformatics, 21(22):4125–4132, 2005.

